# Reward predictability shapes decision making states and exploitative control in marmosets

**DOI:** 10.64898/2026.06.30.734629

**Authors:** Kevin Mastro, Lauren Stanwicks, Erin Schoenbeck, Alex Melain, Matthew Johnson, Bernardo Sabatini, Beth Stevens

## Abstract

Animals navigating dynamic environments must transition between behavioral states dominated by exploitation of known rewards or exploration of alternatives. Understanding this flexibility requires characterizing not only choices and outcomes but also how reward information updates strategy. *Common marmosets* offer a unique combination of genetic tractability and frontal cortical complexity, yet scalable behavioral platforms for studying flexible decision making in this species remain limited. Here, we developed an automated home-cage touchscreen platform that enables voluntary behavioral testing across extended periods without water restriction. We trained marmosets on a series of reversal learning tasks of increasing complexity, culminating in a fully uncued probabilistic two-armed bandit task, and applied reinforcement learning models to examine how reward predictability shapes adaptive choice in marmosets. A Q-learning model with choice perseveration captured trial-by-trial dynamics, with choice variability consistent with stochastic sampling from the fitted policy. A trial-level behavioral state classifier identified distinct and persistent modes of exploitation and exploration that differed in their sensitivity to reward history and their transition structure. Comparing behavioral patterns within probabilistic and deterministic reward contingencies revealed that reward predictability reorganized state occupancy and sharpened error correction within predominately exploitative states, with animals responding to single negative outcomes more rapidly under deterministic feedback. These findings establish both a behavioral and computational framework to dissect the computations and circuits underlying adaptive decision making in marmosets.

## Introduction

Decision making is not a unitary process, but an interplay of dynamic strategies that allow animals to adapt to an ever-changing world. To navigate a non-stationary environment successfully, animals must continuously evaluate whether to exploit known rewards or explore alternatives. The explore-exploit trade-off is shaped by motivation, internal state, and uncertainty, as well as complex interactions between these factors (Daw et al., 2006; Behrens et al., 2007). Disruptions to this balance have been increasingly linked to maladaptive behavioral phenotypes in psychiatric conditions including addiction, anxiety, schizophrenia, and obsessive-compulsive disorder, motivating research into the neural and computational mechanisms that govern flexible decision making (Robbins et al., 2012; Addicott et al., 2017). Harnessing the fluctuations in the explore-exploit tradeoff in the behavior of experimental subjects, rather than treating it as noise, offers a window into the latent strategies that govern moment-to-moment decision making.

At the trial level, choice behavior departs from a purely evidence-based strategy in at least three systematic ways that must be disentangled to characterize choices as exploratory or exploitative. First, spontaneous lapses in performance that appear to reflect disengagement may instead be driven by exploratory processes that are necessary for adaptive flexibility (Ebitz et al., 2019). Second, the average rate of reward shapes response vigor and overall engagement with the task, such that motivational state exerts a tonic influence on the tempo of decision making that is independent of trial-by-trial reward feedback (Niv et al., 2007). Third, perseverative tendencies can dominate behavior in ways that are dissociable from reward-guided learning (Lau and Glimcher, 2005). Identifying these contributions and understanding how they interact requires approaches that can resolve behavioral variability at the level of individual trials across multiple timescales.

Animals transition between behavioral strategies dynamically, but coarse performance metrics obscure this variability, and the factors governing transitions between strategies remain poorly understood. Reinforcement learning (RL) models provide a principled framework for decomposing the latent processes that govern choice in uncertain environments (Sutton and Barto, 1998). Value learning, exploration, and choice perseveration each contribute to choice behavior and can be explicitly separated to identify the underlying mechanisms of behavioral flexibility (Wilson et al., 2014; Gershman, 2018/04/01; Beron et al., 2022). Computational approaches that estimate these parameters at the level of individual trials are therefore necessary to characterize how decision making strategies shift across timescales and task contexts (Ashwood et al., 2022; Bolkan et al., 2022; Mohammadi et al., 2025).

Marmosets are uniquely positioned to bridge the cognitive complexity required to study flexible decision making with the genetic and circuit-level tools needed to investigate its neural basis (Jennings et al., 2016). Marmosets possess a cytoarchitecturally diverse frontal cortex with homologs of human prefrontal regions implicated in value-based choice and behavioral flexibility. Perturbations of distinct prefrontal subregions produce specific and dissociable deficits in reversal learning, value-based choice, and action-outcome contingency tracking, establishing the marmoset as a system for circuit-level dissection of flexible decision making (Rygula et al., 2010; Jackson et al., 2016; Duan et al., 2021b). Building on this foundation, we developed a structured curriculum training protocol delivered by an automated home-cage touchscreen system. This platform enables voluntary participation across extended periods without fluid restriction, progressively guiding animals from simple stimulus-reward associations to a fully uncued probabilistic reversal task. The combination of cognitive complexity, homologous organization of prefrontal cortex, genetic tractability and a scalable home-cage behavioral platform position the marmoset as a pre-clinical model for investigating the circuit mechanisms underlying flexible decision making and their disruption in neuropsychiatric disease.

Here, we characterize how reward predictability shapes the balance between exploitation and exploration in marmosets performing a series of reversal learning tasks voluntarily in their home-cage environment. Voluntary engagement revealed that motivation and reward history make independent contributions to choice behavior, with disengagement occurring predominately following unrewarded trials without disrupting memory of the rewarded action. Applying RL models to trial-by-trial choice sequences, we show that a Q-learning model with choice perseveration captures decision dynamics, and that choice behavior is consistent with a stochastic policy. Using observable trial metrics alone, we identified distinct and persistent behavioral states that differ systematically in their sensitivity to reward information. Finally, comparing behavior under various reward contingencies revealed complementary shifts in exploitation and exploration, demonstrating that the degree of feedback stochasticity shapes decision making in marmosets. Together, these findings demonstrate that voluntary home-cage testing in marmosets can support the rich longitudinal behavioral datasets needed to dissect the computational and circuit mechanisms of flexible decision making, establishing a scalable platform for future mechanistic investigation.

## Results

### Marmosets acquire a visual discrimination task within their home-cage environment and adapt to changes in engagement that modulate performance

To study reward-based decision making, marmosets were trained to interact with a home-cage touchscreen system that enabled repeated behavioral testing over extended periods (**Supplementary Figure 1a**). Individual animals, sequestered into one quadrant of the home-cage, interacted with the apparatus voluntarily during 1-to-3-hour sessions. Marmosets underwent a structured shaping protocol to develop touch behavior such that upon successful completion of each trial, marmosets received sweetened liquid rewards (see *Methods;* **Supplementary Figure 1b**). The animals maintained access to the social housing environment while performing the task and did not require water regulation.

Marmosets (n=10 animals: 7 male/3 female, aged 11-94 months at start of task acquisition) were trained on a visual discrimination task in which two visually distinct stimuli were presented simultaneously at two spatial locations and marmosets indicated their choice by touching the center of the stimuli (**Figure 1a**). For each stimulus pair presented, one was rewarded with probability 1 and the other with probability 0. To test whether marmosets adapt to changing visual cues, we presented marmosets with multiple sets of stimuli within a single session. The contingency for a stimulus pair remained constant for a sequence of trials (known as a block, average block length=33 trials) and a new set of stimuli was introduced at each block boundary. Because both the visual identity of the stimuli and their spatial positions changed at block transitions, animals received a reliable cue signaling a change in stimulus-reward associations. Nonetheless, successful performance required learning which of the two novel stimuli was associated with reward, as the configuration change alone did not indicate which option was better.

**Figure 1.**
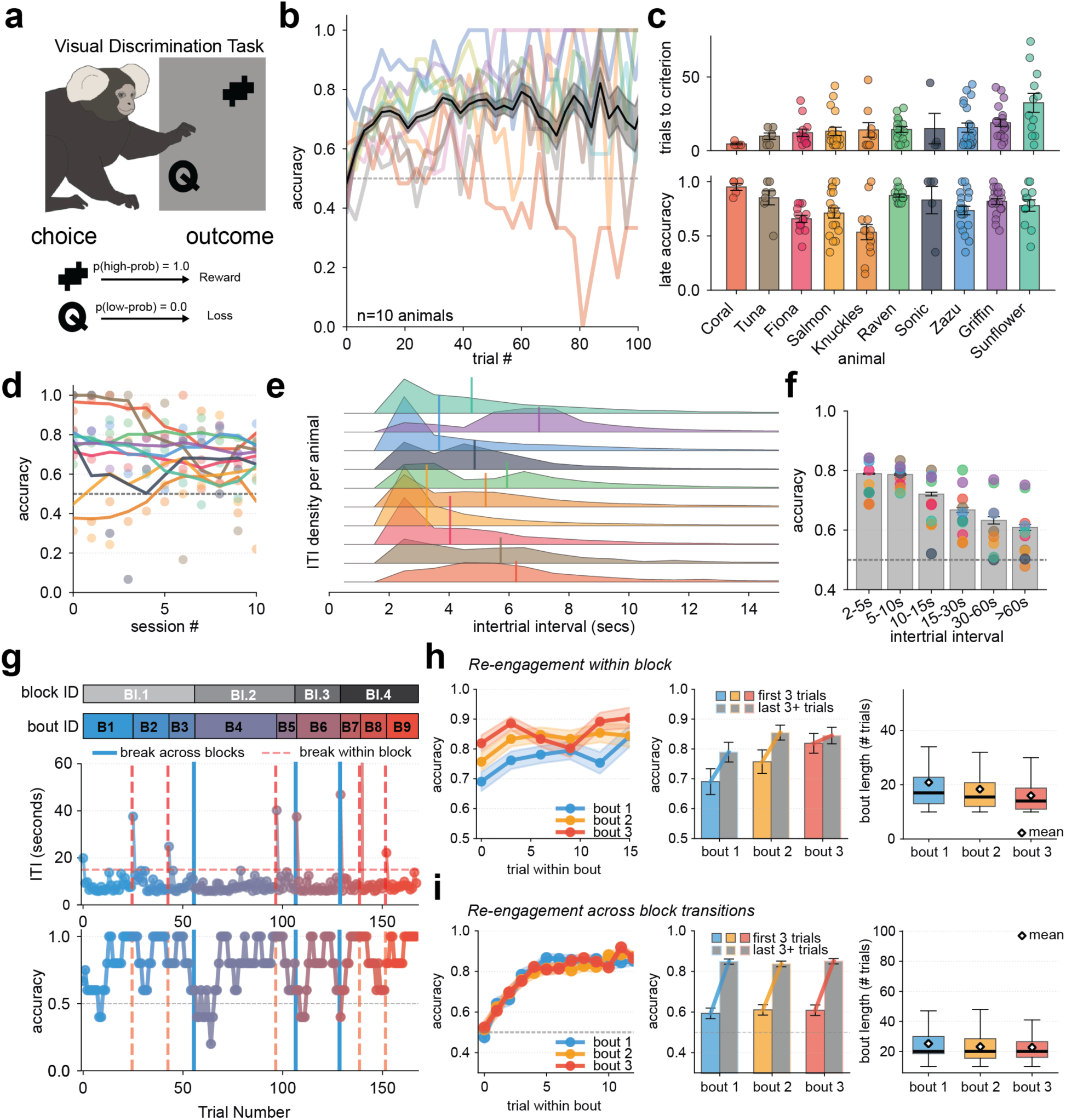
Marmosets acquire deterministic visual discrimination and display improve performance across engagement bouts within and across block transitions. a, Schematic representation of task structure. Marmosets must select one of the two visually and spatially distinct choices by touching a touchscreen. Depending on the choice and task contingencies a juice reward is delivered via a spout. In the visual discrimination task, the probability of reward for selecting one object (the correct choice) is 1 and 0 for other object (the incorrect choice). The identity and location of the rewarded visual object are held constant for a variable length of trials (a block) before changing. b, Probability of selecting the rewarded choice (‘accuracy’) at each trial position on average (black with shaded grey standard error of the mean) and for individual marmosets (colored lines). c, Trials to criterion (top, accuracy >= 0.8 across 5 trials) and the accuracy in the final 10 trials of a block (bottom, “late accuracy”) across all individual marmosets (bars) and sessions (dots). d, Accuracy of selecting the rewarded choice over the course of 10 sessions. Individual dots represent a marmoset’s performance per session, and the colored lines correspond to the performance averaged across multiple days (bin size = 5). Dashed line indicates chance (0.5) e, ITI probability distributions for each marmoset. Medians are marked by the vertical lines. f, Average accuracy for trials binned by the previous ITI for individual animals (colored lines) and aggregated across animals (grey bar plots). g, Representative session showing blocks, bouts, ITIs and performance across one session. Blocks are predefined changes in the visual discrimination rule. Bouts are a series of trials with a maximum ITI of 15 seconds (red vertical dashed line, average 8.8±5.0 bouts per session). Performance is given by the accuracy of selecting the rewarding choice over a 5-trial sliding window. h, Re-engagement within block. Within each bout, line plots show the performance across the first 15 trials (left) followed by bar plots of the aggregate performance at the start and end of the first three bouts (middle) and boxplots of the bout lengths for each bout (right). Bouts shorter than 15 trials contribute all available trials. i, Re-engagement across block transitions. Within each bout, line plots show the performance across the first 15 trials (left) followed by bar plots of the aggregate performance at the start and end of each bout (middle) and boxplots of the bout lengths for each bout (right). Bouts shorter than 15 trials contribute all available trials.

Marmosets successfully learned the cued block transition and stabilized performance over the course of the first 10 trials of each block (**Figure 1b-c**). By the end of each block, marmosets selected the rewarding option in the majority of trials (late accuracy; **Figure 1c**). Performance stabilized across repeated exposure to the task such that all marmosets performed above chance level after 10 sessions of the visual discrimination task (accuracy: 0.664±0.027; Wilcoxon signed-rank vs. 0.5, p=0.002; **Figure 1d**).

Since task initiation was self-motivated and took place within their home environment, marmosets were free to engage or disengage from the task at will throughout each session. On average, individual marmosets had intertrial intervals (ITIs) of 3-7 seconds (**Figure 1e**). During periods of active engagement (trials completed with ITIs < 15 seconds), all marmosets completed bouts of trials that spanned block boundaries, remaining engaged despite changes in task contingencies. During these bouts of engagement across block transition, marmosets actively selected the rewarded stimulus before the change in block (block position -5 to 0; accuracy=0.835±0.003). At the block transition (block position=0), performance decreased to chance levels before recovering over the next ∼5 trials (**Supplementary Figure 1c)**. These results demonstrate that with a high engagement, all marmosets correctly identified and adapted to the demands of a visual discrimination task.

To examine how disengagement affected accuracy, we compared performance across a range of ITI durations. As ITI increased, the accuracy on the subsequent trial declined toward, but remained above, chance (ITI > 60 seconds, mean ITI = 224.5±3.19s accuracy=0.604±0.010; Wilcoxon signed-rank vs. 0.5, p=0.012; **Figure 1f**). To assess how periods of disengagement impacted decision making, we defined bouts of behavior as consecutive trials with ITIs < 15 s with a new bout beginning after longer periods of inactivity (ITI ≥ 15 s; corresponding to the 90^th^ percentile of ITI distribution).

Bouts of behavior occurred throughout each session (8.8±5.0 bouts per session), arising either within a block or across block transitions (**Figure 1g**), and the termination of a bout was more frequent following an unrewarded trial (**Supplementary Figure 1d-e)**. For bouts occurring within a block, accuracy at re-engagement remained above chance across successive bouts while bout length remained constant (Bout 1: 18.9 trials, Bout 2: 18.1 trials, Bout 3: 16.0 trials), indicating that marmosets retained memory of the rewarded action across periods of disengagement (**Figure 1h**). At the start of a new block, marmosets frequently experienced an unrewarded trial that led to a prolonged period of disengagement. Upon re-engagement, marmosets identified the rewarded action and achieved criterion (accuracy > 0.75) within 5.3±0.1 trials and completed 20 or more trials (**Figure 1i**). These findings demonstrate that marmosets adapted to changes in stimulus-reward relationships even across periods of disengagement, retaining not only memory of the rewarded action but an inference about which action to pursue upon re-engagement.

### Visual discrimination task with reversal reveals reward-dependent shifts in behavior across both cued and uncued transitions

Changes in action-outcome relationships in the visual discrimination task were signaled by simultaneous changes in stimulus appearance and spatial location, providing an observable cue at each block transition. To probe the degree to which decision making depends on reward history rather than stimulus changes, we introduced reward-probability reversals within each stimulus pair such that the identity of the rewarded and unrewarded stimuli flipped (**Figure 2a**). Critically, the visual stimuli and their spatial positions remained constant across these reversals, eliminating overt cues that task contingencies had changed. Within each session, marmosets experienced both cued transitions (changes in visual stimuli and spatial configuration) and uncued reversals (changes in reward contingency only), allowing direct comparisons between the two. Cued and uncued transitions therefore exercise distinct processes: cued transitions prompt the animal to seek the newly rewarded stimulus, whereas uncued reversals require the animal to infer that contingencies have changed from reward outcomes alone before updating its strategy.

**Figure 2.**
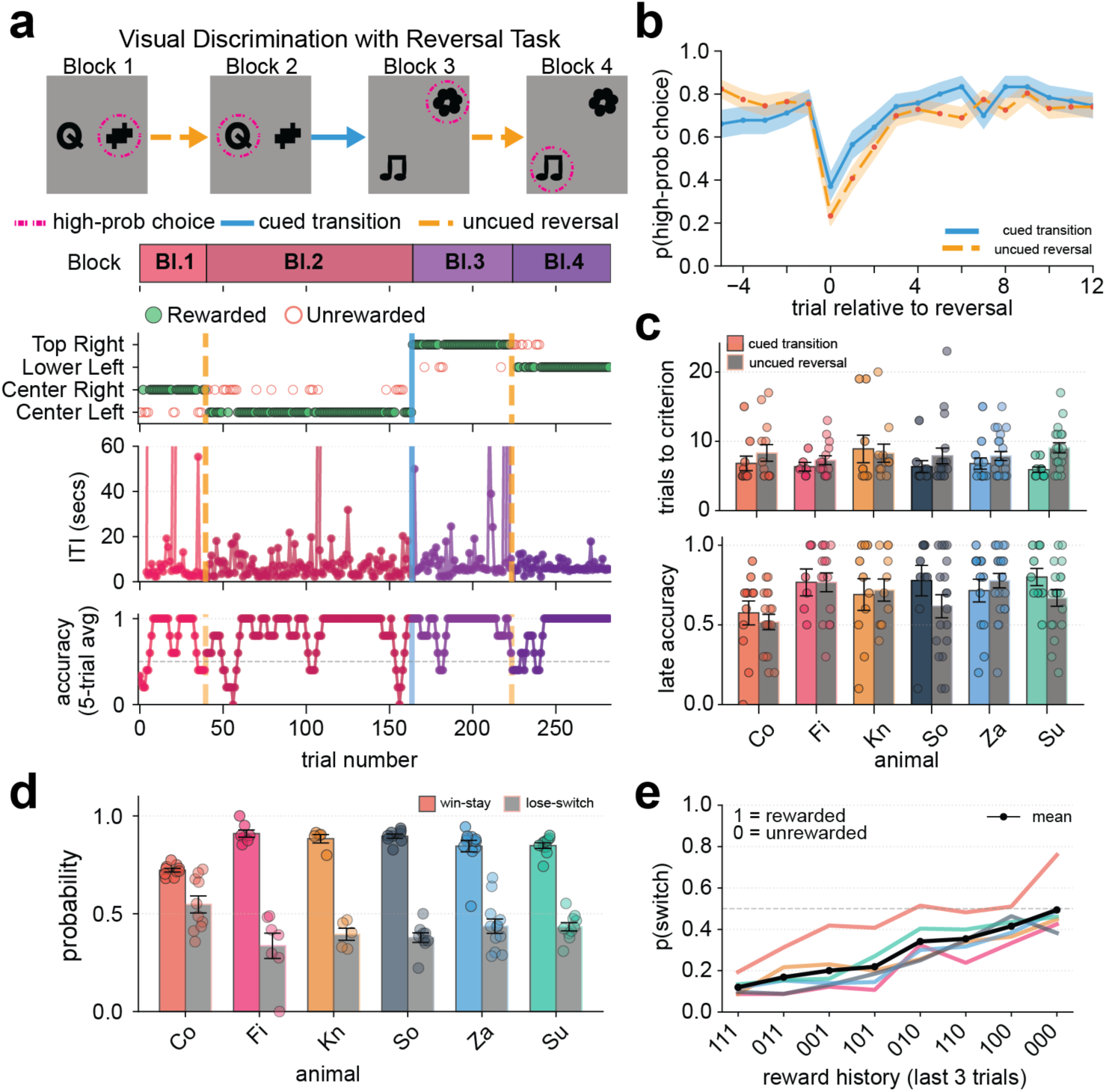
Marmosets update switching behavior based on reward history. **a**, Schematic representation of task structure and example showing blocks, ITIs and performance across one session. In the visual discrimination with a reversal task, the probability of reward for selecting one object (the correct choice) is 1 and 0 for other object (the incorrect choice). The identity and location of the rewarded visual object are held constant for a variable length of trials (a block) before reversing the reward contingencies for the two stimuli. Following a reversal block, the next block will be a cued transition where two new visual stimuli will appear in two new locations on the touchscreen. Blocks are predefined changes in the visual discrimination or reversal rule; vertical lines indicate whether change is a “cued transition” (solid blue) or an uncued reversal (dashed orange). Performance is given by the accuracy of selecting the high probability choice over a 5-trial sliding window. **b**, Probability of selecting the high probability choice (‘high-prob choice’) at each trial position for either cued transition (blue solid line) or uncued reversal (orange dashed). Shaded error bars indicate standard error of mean. **c**, Trials to criterion (top, accuracy >= 0.8 across 5 trials) and the accuracy in the final 10 trials of a block (bottom, “late accuracy”) across all individual marmosets (bars) and sessions (dots). **d**, Probability of repeating after a reward (win-stay) or switching after a loss (loss-switch) across all individual marmosets (bars) and sessions (dots). **e**, Probability of switching conditioned on the three previous outcomes (rewarded = 1, unrewarded = 0) where the most recent outcome is the final number of the sequence, averaged across all marmosets (black line) superimposed on top of individual marmosets (colored lines).

Across reversals, marmosets displayed three features of reward-based learning. First, the trials to criterion to new stimulus-reward contingencies was similar between cued and uncued transitions. Marmosets reached 75% accuracy within 7.4±1.4 trials following cued transitions and 6.7±0.3 trials following uncued reversals, and these did not differ significantly (**Figure 2b-c**). Late-block accuracy was comparable across transition types (Cued: 0.749±0.017, Uncued: 0.677±0.040; Wilcoxon signed-rank, p=0.44; **Figure 2c**). Second, individual animals varied in their sensitivity to reward feedback, with some marmosets adapting rapidly to uncued reversals while others required extended experience (**Figure 2c**). Importantly, the informational content of unrewarded trials differed between cued and uncued contexts: in the visual discrimination task, a loss indicated an incorrect choice, whereas during uncued reversals, a loss at a previously rewarded choice signaled that task contingencies had changed. To quantify the dynamics of choice updating, we examined the probability of switching choices following rewarded and unrewarded trials. Consistent with reward-based learning, marmosets were more likely to repeat rewarded choices (win-stay: 0.852±0.026) while switching more frequently following unrewarded choices (lose-switch: 0.401±0.023; **Figure 2d**).

While win-stay and lose-switch probabilities capture single-trial reward sensitivity, the third feature addresses how reward history accumulates across multiple trials. Animals did not immediately and stably switch to the newly rewarded option after a single unrewarded trial. Instead, choice behavior shifted gradually across trials, consistent with perseverative tendencies slowing the transition to the newly rewarded option. To quantify how reward history influenced switching, we binned trials based on the three preceding outcomes (Rewarded=1, Unrewarded=0, with the most recent trial in the third position) and calculated the probability of switching on the subsequent trial. A trial history of “110” (two rewards followed by one omitted reward) yielded an average switching probability of 0.350±0.033, whereas “000” (three consecutive omitted rewards) yielded a switching probability of 0.454±0.027 (**Figure 2e**). In contrast, three consecutive rewarded trials produced the lowest switching probability (0.126±0.017), demonstrating that reward history drives behavioral adaptation with respect to reward history. These results indicate that marmosets’ behavioral adaptation is guided by accumulated reward history even in the absence of explicit task cues, motivating the development of a fully uncued probabilistic reversal task to isolate reward-based learning from stimulus-driven updating.

### Marmosets adapt to probabilistic reward contingencies during uncued reversals in a two-armed bandit task

The visual discrimination with reversals task demonstrates that marmosets can adapt to changing reward contingencies even without external cues, but each session still contains cued transitions that could influence overall strategy. These tasks were completed under deterministic reward probabilities for which the simplest strategy (win-stay, lose-switch) is maximally effective. To isolate purely reward-based learning, we designed a two-armed bandit task (2-ABT) in which all transitions were uncued and reward contingencies were probabilistic, as used previously in rodents (Beron et al., 2022; Mastro et al., 2025). In the 2-ABT, marmosets chose between two spatially distinct options (e.g. left or right touchscreen locations) that yielded probabilistic rewards. Within each block, one location yielded reward with probability 0.8 and the other with probability 0.2. At block boundaries, these probabilities reversed without any change in visual appearance or spatial locations, requiring animals to infer reversals solely from the pattern of rewards and reward omissions over a series of choices (**Figure 3a**).

**Figure 3.**
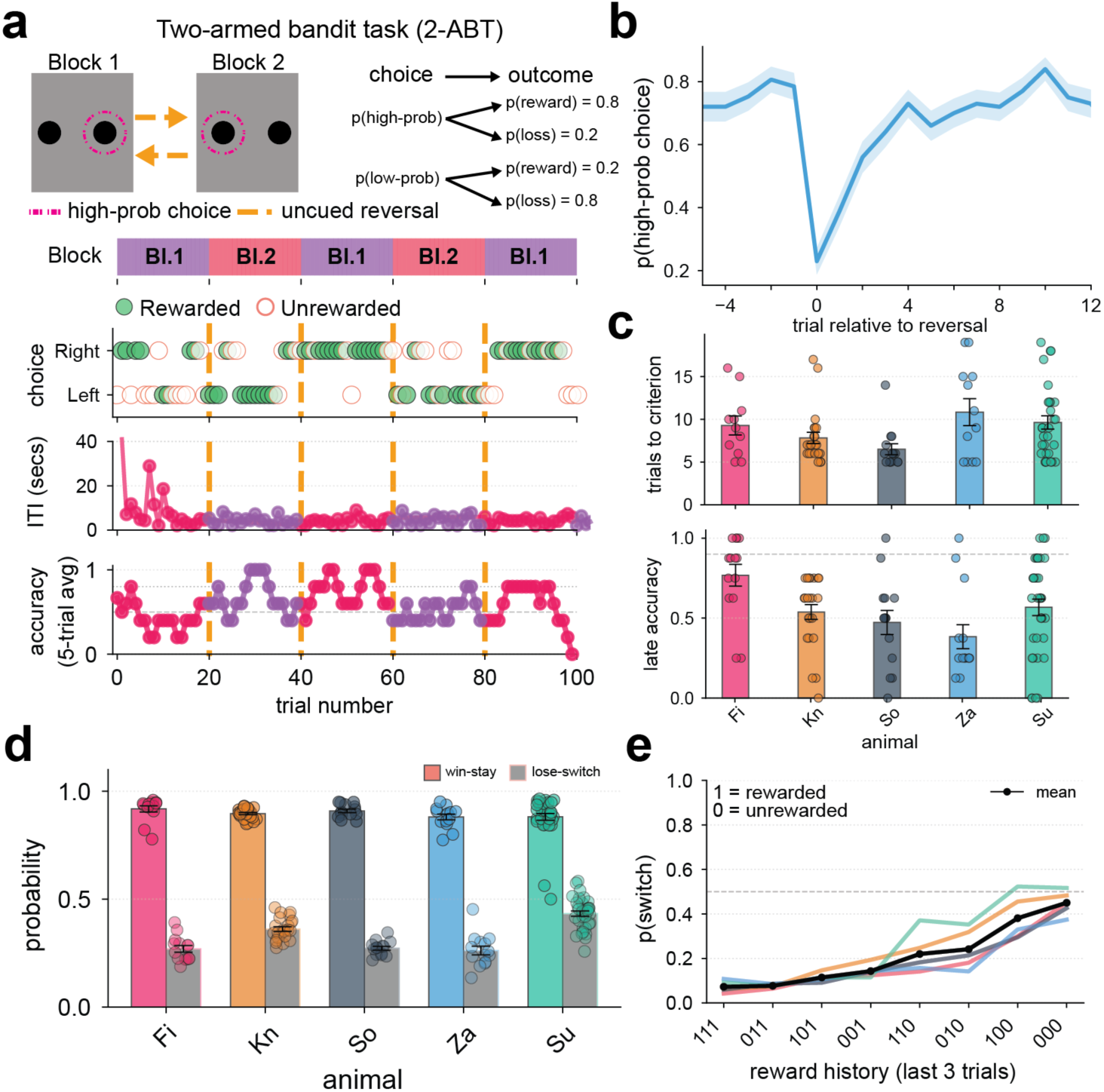
Marmosets acquire a 2-armed bandit task. **a**, Schematic representation of task structure and example showing blocks, ITIs and performance across one session. In the 2-armed bandit task, the probability of reward for selecting one object (high probability choice) is 0.8 and 0.2 for other object (low probability choice). The identity and location of the rewarded visual object are held constant for a variable length of trials (a block) before reversing the reward contingencies for the two stimuli (uncued reversal, vertical dashed line) Performance is given by the accuracy of selecting the high probability choice over a 5-trial sliding window. **b**, Probability of selecting the high probability choice (‘high-prob choice’) at each trial position across an uncued reversal where shaded error bars indicate standard error of mean. **c**, Trials to criterion (top, accuracy >= 0.8 across 5 trials) and the accuracy in the final 10 trials of a block (bottom, “late accuracy”) across all individual marmosets (bars) and sessions (dots). **d**, Probability of repeating after a reward (win-stay) or switching after a loss (loss-switch) across all individual marmosets (bars) and sessions (dots). **e**, Probability of switching based on the three previous outcomes (rewarded = 1, unrewarded = 0) averaged across all marmosets (black line) superimposed on top of individual marmoset (colored lines).

Under probabilistic reward contingencies (80-20), marmosets successfully tracked reward probabilities and adapted to reversals across blocks. Five animals completed 20±4 sessions of the probabilistic 2-ABT, performing 367±14 trials per session. The probability of selecting the high-probability choice stabilizes by the end of each block (trials -5 to -1: accuracy=73.6±0.5%, **Figure 3b**). At the reversal (block position=0), marmosets initially selected the previously high-probability choice and required nearly 10 trials to achieve stable performance (**Figure 3b-c**). Across all blocks (n=1,779 across 100 sessions), 98.2% reached criterion (p(high-probability choice)>=0.75) performance within 6.8±0.2 trials of the reversal, consistent with the interpretation that animals reliably adapted to changes in reward contingencies.

Marmosets showed the same characteristic win-stay/lose-switch asymmetry in the 2-ABT, but with a pattern consistent with the increased uncertainty of probabilistic reward contingencies. Comparing across animals run on both tasks, win-stay probability was higher in the 2-ABT than in the visual discrimination task with reversals (2-ABT: 0.903±0.006 vs. VDT: 0.877±0.013; t(4)=−3.851, p=0.018), while lose-switch probability did not differ significantly between tasks (2-ABT: 0.327±0.035 vs. VDT: 0.396±0.019; t(4)=2.281, p=0.085; **Figure 3d**). These results are consistent with single unrewarded trials being less informative when rewards are inherently variable. Nonetheless, switch probability increased in a graded manner with the number of recent unrewarded outcomes, rising from 0.073±0.008 following three consecutive rewards (‘111’) to 0.450±0.027 following three consecutive omissions (‘000’; **Figure 3e**), confirming that accumulated reward history guides behavioral adaptation in a graded manner even under probabilistic contingencies.

### Reinforcement learning models with choice perseveration capture trial-by-trial decision dynamics

To understand the computational mechanisms underlying adaptive behavior in the 2-ABT, we fit a variety of reinforcement learning (RL) models to individual choice sequences. In these models, estimates of action values (i.e., Q-values) for each pair of visual stimuli are stored, and updated on each trial based on prediction errors, defined as the difference between received and expected rewards (Sutton and Barto, 1998). We compared models that differed in their assumptions about how past choices and rewards influence current decisions.

Model selection was based on the ability for the model to predict trial-by-trial performance as defined in the following analyses: probability of selecting the high-prob choice across block transitions, probability of switching across block positions and the comparison between reward history-conditioned switching probabilities for the marmoset vs the model (**Supplementary Figure 2a-d**). After comparing Q-learning models (see **Methods**), we selected a Q-learning model with a choice kernel as the working model for subsequent analyses based on parameter stability and interpretability. The model captures trial-by-trial fluctuations in choice behavior, correctly predicting choices on 81.0 ± 1.4% of held-out test trials and accurately reproducing the temporal dynamics of log-odds of choosing right versus left (**Supplementary Figure 2e**).

The Q-learning model with a choice kernel successfully captures the tendency to repeat recent choices independent of their reward outcomes (**Figure 4a**). In this model, Q-values for the chosen option are updated according to:

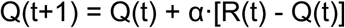

**Figure 4:**
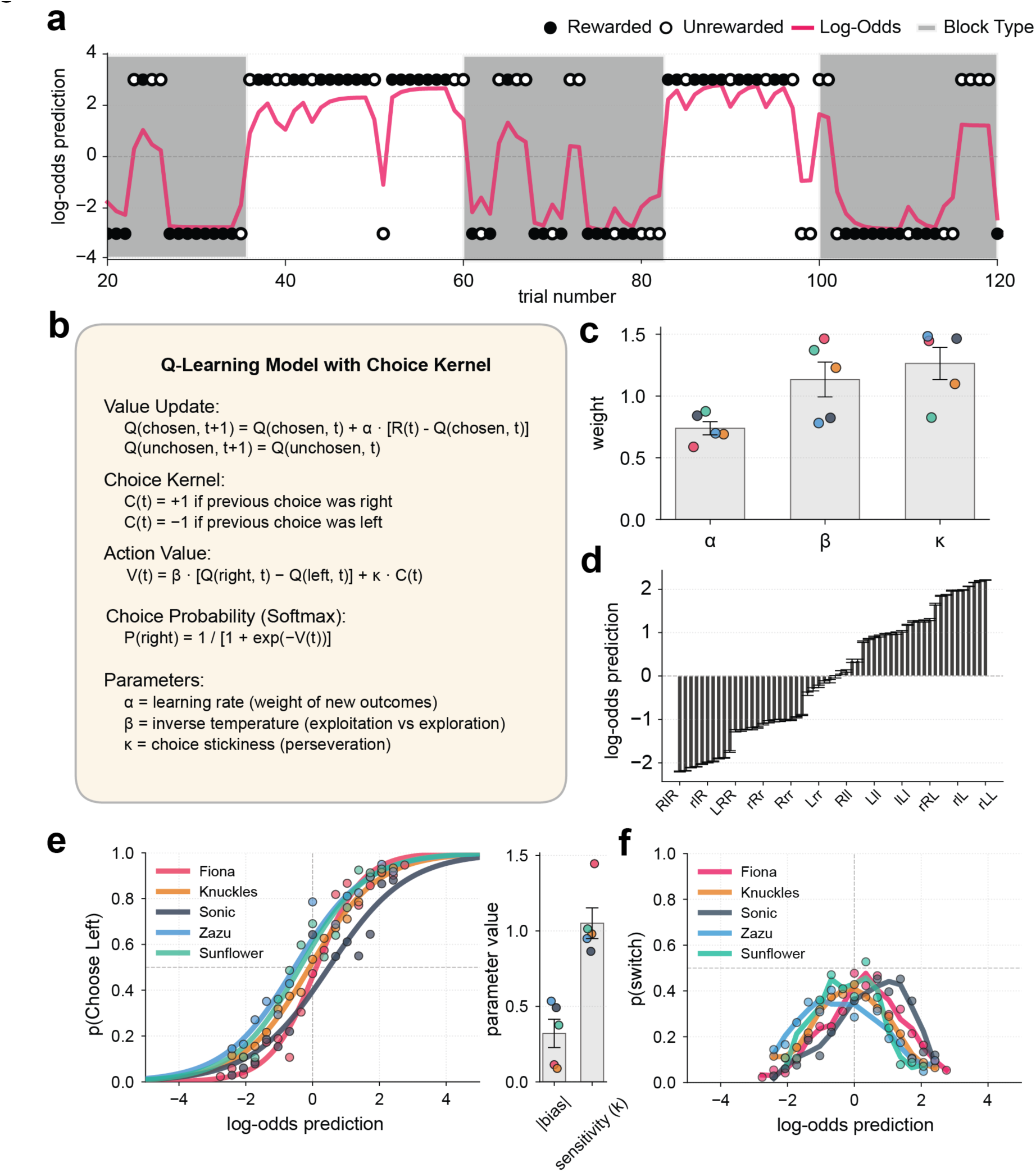
Q-learning model with choice kernel captures trial-by-trial dynamics of the 2-armed bandit performance and captures differences across individuals. **a**, Example session depicting the choice-outcome (Choice: Top /Bottom, Outcome: Rewarded = filled, unrewarded = unfilled) across a subset of trials. Shaded regions indicate that the bottom is the high probability choice. Overlaid is the updated q-values across the session (magenta line). **b**, Q-learning Model with Choice kernel functions and policy (Softmax). **c**, Bar plot depicting the log-odds predictions binned by the action-outcome histories **d**, Average of model parameters (grays bars) overlaid with individual animal means (dots) across alpha (learning rate), beta (inverse temperature) and kappa (sticky term). **e**, Probability of choosing the left choice binned across the log-odds prediction for individual marmosets (dots) and the sigmoid function fits across values (solid lines) (left). Bar plot of the average (bars) and individual (dots) values of the bias and sensitivity derived from the sigmoid fit parameters. **f**, Probability of switching binned across the log-odds prediction for individual marmosets (dots) and the sigmoid function fits across values (solid lines)

where α represents the learning rate and R(t) is the reward received (1 for rewarded, 0 for unrewarded trials). The choice kernel, C(t), tracks whether the animal chose the same option as on the previous trial (C(t)=1) or switched (C(t)=0). The action value on each trial combines learned value and choice history:

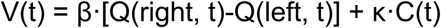

where β represents inverse temperature (the degree to which choices favor higher-value options) and κ represents choice perseveration (the tendency to repeat recent actions beyond what learned value alone predicts) (**Figure 4a-b**).

The behavior of each marmoset is best described by models with different parameter values (**Figure 4c**). Learning rates (α) averaged 0.746±0.073 across animals, indicating that animals weighted recent outcomes heavily while retaining some influence of older reward history. At β=1 choices are made in proportion to value differences; values greater than 1 indicate increasing selectivity for the higher-value option, while values less than 1 and approaching 0 produce near-random choice. Inverse temperature (β) values of 1.115±0.155 are consistent with a stochastic rather than deterministic choice policy, by which animals favor higher-value options but sample lower-value options probabilistically. Choice perseveration (κ) of 1.298±0.147 is consistent with a tendency to repeat recent choices beyond what learned value predicts alone.

To validate that the model captures meaningful structure in behavior, we examined whether the model’s internal estimates of choice probability predict actual choices. On each trial, the model computes a log-odds score quantifying the probability of selecting the right versus left option based on learned Q-values and choice history. First, to characterize how action-outcome history maps onto these predictions, we encoded the three trials preceding each choice as a sequence of symbols denoting action and outcome: choice was denoted with either R (right) or L (left) while the outcome was denoted by either uppercase (rewarded) or lower case (unrewarded). These sequences were compared against the model’s internal predictions of choice for the next action (log-odds prediction; **Figure 4d**). As expected, the model’s log-odds predictions tracked action-outcome history in an interpretable manner: positive log-odds are dominated by recently rewarded choices on the left side (e.g. ’LLL’, ’rLL’), whereas negative log-odds are dominated by histories with recent rewards on the right side (e.g. ’RRR’, ’RlR’). This is consistent with the model’s internal estimates reflecting both direction and the recency of rewards in a behaviorally meaningful way. Second, we binned trials according to these log-odds predictions and compared the model’s predicted choice probability to the empirical choice probability within each bin. Across the full range of preferences, from strong right preference (log-odds < -2) to strong left preference (log-odds > +2), model predictions and observed behavior align closely (**Figure 4e**, mean r = 0.951 ± 0.025; all p < 0.001) consistent with the model’s log-odds estimates tracking animals’ trial-by-trial choices.

Finally, we examined whether the model captures the dynamics of switching behavior. Empirical switch rates show a characteristic inverted U-shaped relationship with log-odds of left/right choice. Animals switch most frequently when preference is near zero and switch least when strongly preferring one option, a pattern expected under a softmax decision policy, in which choice stochasticity is highest when action values are similar (**Figure 4f**). The Q-learning model with choice kernel reproduces this pattern, consistent with a stochastic policy in which choice variability is highest when the decision variable approaches zero. Together, these results are consistent with a simple RL framework in which value learning from reward feedback combined with choice perseveration provides a parsimonious account of marmoset decision making in the 2-ABT. The fitted Q-values from this model provide trial-by-trial estimates of learned value that can be used to probe whether and how animals track reward information across different behavioral contexts.

### Marmosets Q-value adherence is consistent with a stochastic policy, but asymmetrically modulated by reward outcome

To assess the degree to which animals’ choices are consistent with the choice value estimates of the Q-learning model, we identified on each trial whether the animal selected the option with the higher estimated value (Q-adherence) or the lower estimated value (Q-violation). Under the fitted model, choice probability follows a softmax policy in which selection depends on the decision variable V(t), such that choices become more stochastic as value estimates converge and choice perseveration provides weaker directional guidance. Representative sessions illustrated the behavioral contrast between low and high violation rate sessions. In low violation sessions, choices track the dominant Q-value closely and switch reliably at block transitions. In high violation sessions, animals frequently depart from the model-preferred option and persist on the non-preferred side even as Q-values diverge (**Figure 5a**). Across all animals and sessions, marmosets adhere to their Q-values on 71.7% of trials, with violation rates consistent across individuals (29.6±1.7%; **Figure 5b; Supplementary Figure 3a**).

**Figure 5.**
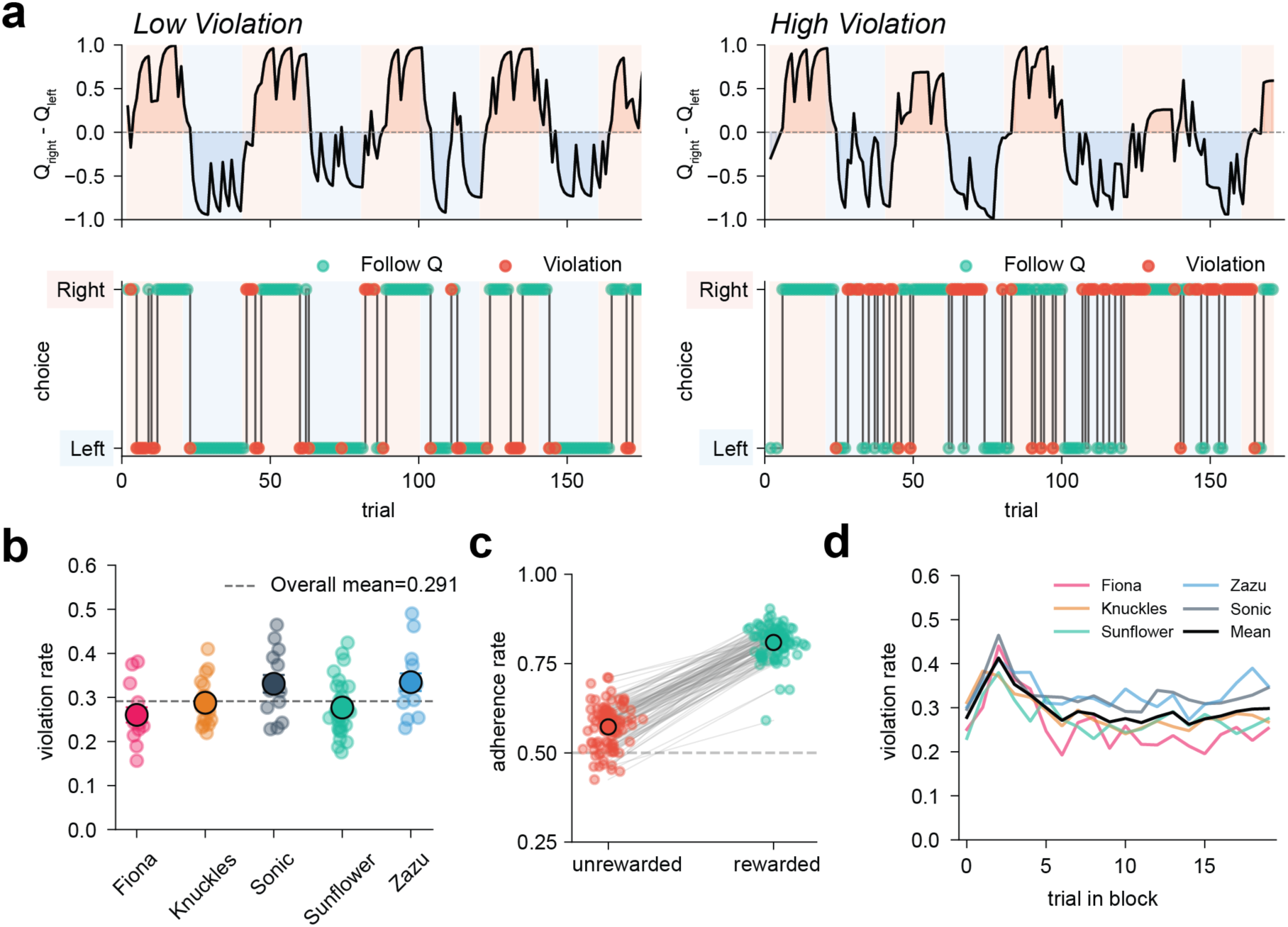
Marmosets adhere to learned Q-values on majority of trials, with violations occurring preferentially under conditions of uncertainty. **a**, Representative sessions from a single animal (Fiona) illustrating low (left) and high (right) violation rate sessions. (top) Trial-by-trial difference in Q-values between the right and left options (Q_right_ − Q_left_), computed from a Q-learning model with choice perseveration term. Background shading indicates which side was the high-probability option (red: right high-prob; blue: left high-prob). (bottom) Choice sequence (right/left) with individual trials colored by whether the animal adhered (green) or violated (red) its Q-values. **b**, Violation rate by animal. Each dot represents one session; large, filled circles indicate the session mean. Dashed line indicates the overall mean violation rate across all animals and sessions (0.291). **c**, Adherence rate on unrewarded versus rewarded trials. Large, filled circles indicate the mean across all sessions and animals while gray lines connect matched sessions (dots). Animals were more likely to adhere to Q-values following a rewarded outcome (p < 0.0001, paired t-test). **d**, Violation rate as a function of trial position within a block. Colored lines show individual animal means; black line shows the mean across all sessions and animals.

To determine whether this violation rate reflects stochastic sampling from the fitted policy or systematic departures from value-guided choice, we compared observed choice consistency against the theoretical ceiling predicted by the model’s stochastic policy. For a softmax policy, the probability that a single draw from the policy produces the argmax choice or, in other words, the probability that two independent draws from the policy agree is given by a theoretical quantity we refer to as the policy-predicted adherence rate (see **Methods**). Across animals, observed P(Q-adherence) did not differ from the policy-predicted adherence rate (observed: 0.704 ± 0.017, policy-predicted: 0.722 ± 0.013; t(4) = −0.871, p = 0.433; **Supplementary Figure 3b**), indicating that the aggregate violation rate is consistent with stochastic sampling from the fitted policy rather than systematic departures from value-guided choice.

Although the aggregate violation rate is policy-consistent, examining Q-value adherence across reward outcomes revealed that the single fitted choice perseveration parameter κ does not uniformly describe choice behavior. The probability of repeating the previous choice was substantially higher after rewarded trials than after unrewarded trials (rewarded: 0.798 ± 0.063, unrewarded: 0.573 ± 0.079; t(4) = 36.675, p < 0.00001, Cohen’s d = 3.887; **Figure 5c**), and this asymmetry exceeded what the single fitted κ predicts. After rewards, the policy-predicted adherence rate sat above the model prediction across all levels of value certainty while it sat below after losses (**Supplementary Figure 3c**). Fitting a model with separate choice perseveration parameters for post-reward and post-loss trials confirmed this asymmetry: κ_win_ = 2.125 ± 0.076 was substantially larger than κ_loss_ = 0.774 ± 0.163 across all five animals (t(4) = 7.079, p = 0.0021), with the asymmetric model providing substantially better fit than the single-κ model (mean ΔBIC = −222.0). These results indicate that reward outcome modulates the strength of choice repetition beyond what a single choice perseveration parameter captures, with animals showing stronger win-stay and lose-switch tendencies than the symmetric policy assumes.

The across-trial profile of violations within blocks similarly tracks the model’s expected certainty. Violation rate was highest immediately after block transitions where Q-values had not yet updated to reflect the reversed reward contingency and thus provided weak or misleading guidance. After this period, the violation rate decayed toward the session mean within approximately five trials as animals accumulated evidence for the new contingency (**Figure 5d**; **Supplementary Figure 3c**). This time course mirrored the learning dynamics observed in the reversal-aligned performance curves (**Figure 3b**), suggesting that the same updating process that drives accuracy recovery also governs the recovery of Q-value adherence after reversals.

At the population level, violations are directionally unbiased such that the proportion directed toward the left versus right option did not differ from 50% (p=0.093), but individual animals show consistent spatial tendencies in their Q-violation behavior (**Supplementary Figure 3d**). As expected from trial-by-trial reliance on an estimate choice value, trials after a Q-violation have both reduced reward probability and reduced probability of Q-value adherence on the subsequent trial compared to when an animal adhered to their Q-values (**Supplementary Figure 3e-f**; both p < 0.0001, paired t-test), indicating that violations initiate brief periods of increased behavioral variability rather than representing isolated bouts of task rule departures. At the session level, Q-violation rate is negatively correlated with accuracy for Fiona (r=-0.61, p=0.021), Sonic (r=-0.62, p=0.017) and Sunflower (r=-0.77, p < 0.001), but is not significantly related to performance for Knuckles or Zazu (**Supplementary Figure 3g**), suggesting that the behavioral consequences of violating Q-value guidance vary across individuals.

The Q-value analysis establishes that aggregate violations are consistent with the model’s stochastic policy and that reward outcome modulates choice repetition beyond what a single choice perseveration parameter captures. However, the Q-value framework alone cannot determine whether trial-by-trial variability reflects genuine uncertainty-driven exploration or spatial perseveration independent of learned value. We therefore examined whether distinct behavioral states could be identified from observable behavior alone, and whether these states differ systematically in their relationship to learned value and reward history.

### Unsupervised clustering delineates distinction between exploration, exploitation and bias

Departures from reward-guided choice occur preferentially under conditions of value uncertainty but do not in themselves distinguish animals actively exploring to resolve that uncertainty from animals perseverating on a spatial preference independent of learned value. To identify whether distinct behavioral states could be recovered from observable choice metrics alone, we applied unsupervised K-Means clustering and a Gaussian Hidden Markov Model to four behavioral features computed over a five-trial sliding window: absolute deviation from selecting the high-probability option, rolling average of accuracy, spatial choice deviation, and rolling switch rate, metrics derived from behavior and task structure without reference to the RL model.

K-Means clustering identified four behavioral clusters, with the number of clusters selected by silhouette coefficient, which peaked at K=4 while both statistical criteria (Akaike Information Criterion, AIC and Bayesian Information Criterion, BIC) showed a modest elbow at K=4 (**Supplementary Figure 4a**). Based on their cluster profiles, these four clusters corresponded to Exploitation, Exploration, Left Spatial Bias, and Right Spatial Bias (**Figure 6a)**. Cluster profiles revealed the behavioral characteristics of each state: the Exploitation cluster had high rolling accuracy (0.786) and low switch rate (0.076), the Exploration cluster had moderate accuracy (0.461) and high switch rate (0.420), and both bias clusters were defined by strongly lateralized signed deviation (Left Bias: −0.639, Right Bias: +0.643) with moderate accuracy and low switch rates (**Supplementary Figure 4b**). Principal component analysis of the four features reveals the structure of the clustering across three components (**Figure 6b**). PC1 (37.9% of variance) separated Exploitation from Exploration and Bias states along a performance axis. PC2 (28.1% of variance) further separated Bias states from exploration states within the low-PC1 region. PC3 (25.0% of variance) separated left from right spatial bias along a lateralization axis, orthogonal to the performance and switching dimensions. The majority of trials were spent in either Exploitation or Exploration states and the remaining trials were spent equally across the Right and Left Bias states (**Figure 6c**).

**Figure 6.**
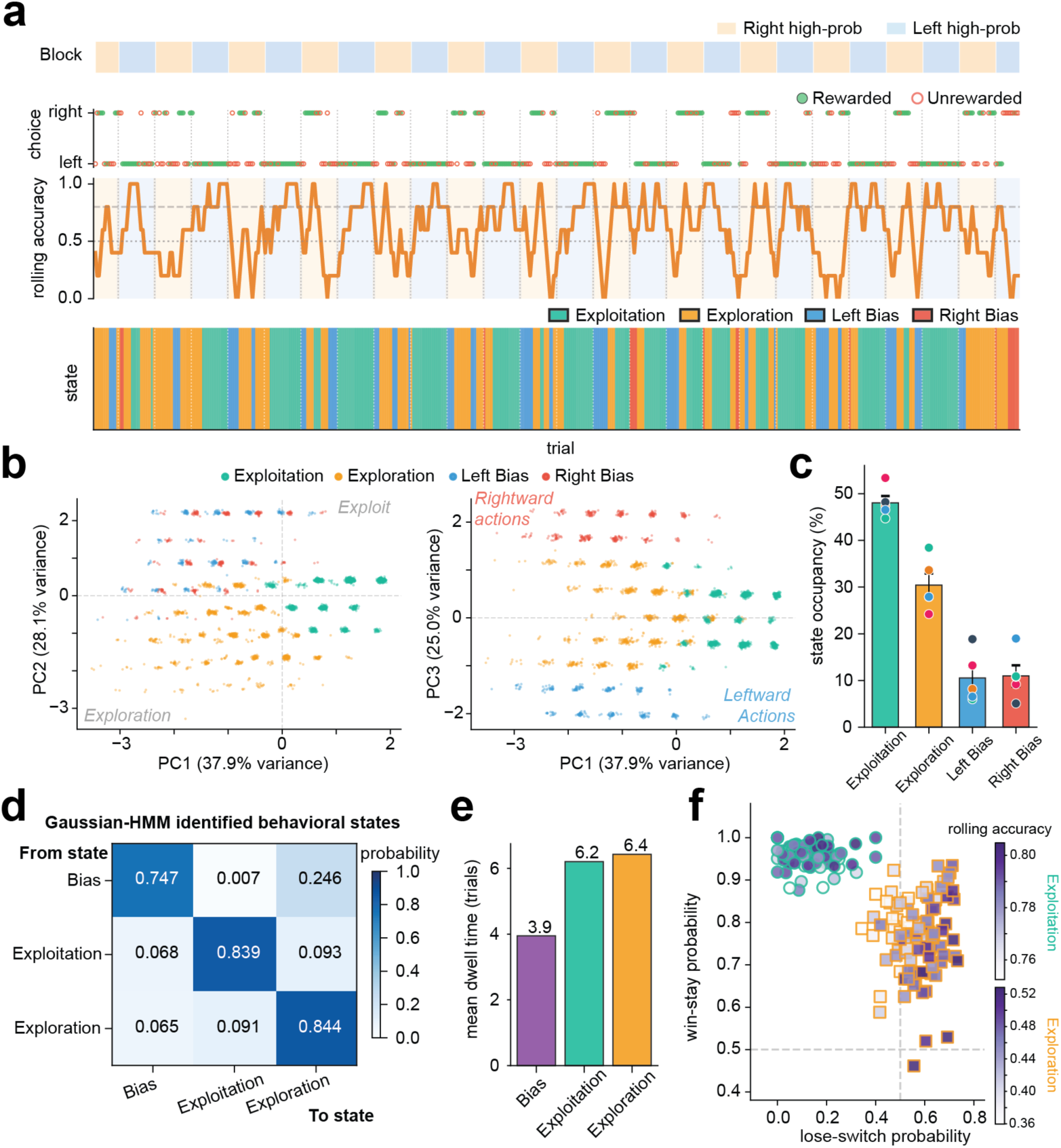
Unsupervised clustering recovers distinct behavioral states from observable choice metrics. **a**, Representative sessions from a single animal (Knuckles) showing block structure (top), rewarded and unrewarded choices and rolling accuracy computed over a five-trial window (middle), and trial-by-trial K-means state classification (bottom). Colors correspond to the four states defined in the legend. **b**, Principal component projections of the behavioral feature space colored by K-Means cluster identity. Left: PC1 (37.9% variance) versus PC2 (28.1% variance). PC1 separates Exploit from Exploration and Bias states along a performance axis; PC2 separates Bias from Exploration within the low-PC1 region. Right: PC1 versus PC3 (25.0% variance). PC3 separates left from right spatial bias along a lateralization axis. Small Gaussian jitter (σ=0.03) added for visualization. Colors indicate cluster identity (Exploit: teal; Exploration: orange; Left Bias: blue; Right Bias: red). **c**, State occupancy (% of trials) per behavioral state. Bars indicate the mean across animals; dots indicate individual animal means (n=5). **d**, Transition probability matrix from a Gaussian Hidden Markov Model fit to behavioral features. Row-normalized to give the conditional probability of transitioning from each state (rows) to each state (columns). Direct transitions from Bias to Exploitation were almost never observed (p=0.007). **e**, Mean dwell time per HMM state, derived analytically from self-transition probabilities. **f**, Win-stay versus lose-switch probability for Exploitation and Exploration trials. Each point represents one session colored by rolling accuracy within state (Exploitation: teal-to-purple; Exploration: orange-to-purple). Circles indicate Exploitation trials; squares indicate Exploration trials. **a-f.** N=5 animals, n=100 sessions total. All classification and clustering analyses were performed on held-out sessions not used for threshold derivation.

To delineate the sequential structure between states, a Gaussian Hidden Markov Model (HMM) fit to the same features independently recovered three states: Bias, Exploitation, and Exploration. For the HMM, the signed choice deviation did not improve state-recovery and therefore only the absolute was used. The HMM transition matrix revealed that direct transition from Bias to Exploitation states was almost never observed (p(Bias to Exploitation)=0.007) with state changes instead routed through Exploration as an intermediate (**Figure 6d**). Mean dwell times derived analytically from self-transition probabilities were 6.2 trials for Exploitation, 6.4 trials for Exploration, and 3.9 trials for Bias (**Figure 6e**), indicating that bias episodes are more transient than periods of exploitation or exploration. HMM emission profiles confirmed the behavioral interpretation of each state, with Bias characterized by high spatial deviation and moderate accuracy, Exploitation by high accuracy and near-zero switch rate, and Exploration by elevated switch rate and intermediate accuracy (**Supplementary Figure 4c**). This distinction was also evident in reward sensitivity: per-session win-stay and lose-switch probabilities, computed separately for Exploitation and Exploration trials within each session, occupied separable regions (**Figure 6f**). Sessions showed high win-stay and low lose-switch probability during Exploitation, while the same sessions showed substantially higher lose-switch probability and more variable win-stay during Exploration, indicating that the two states are governed by distinct combinations of reward sensitivity rather than a shared asymmetry.

Agreement between HMM-decoded states and K-Means clusters on held-out test sessions was moderate (Adjusted Rand Index (ARI): 3-state ARI=0.487; 4-state post-hoc ARI=0.474), consistent with both methods recovering similar large-scale behavioral structure from choice data alone. Notably, agreement was highest for the bias state (confusion matrix precision=1.0), indicating that spatial perseveration was robustly identified by both methods, while disagreement was concentrated at the exploitation/exploration boundary (HMM-exploitation trials classified as exploration by K-Means: 39.2%), consistent with a gradual rather than discrete transition between exploitation and exploration. Together, these results indicate that observable choice behavior contains sufficient structure to recover distinct and interpretable decision making states without reference to a computational model.

### A behavioral state framework reveals distinct modes of reward-guided decision making in marmosets

The three-state solution recovered by unsupervised clustering and the modest alignment between the identified states raises two important questions: 1) whether finer-grained structure exists within exploration and exploitation and 2) how the discrete contingency changes imposed by block reversals are represented within this framework. To address this, we defined a behavioral taxonomy in which boundaries and transition structure reflect empirically identified discontinuities in behavioral space (**Figure 7a**). Like the unsupervised clustering, we used the same set of observable choice metrics and delineated additional structure to the decision making behavior. Within this taxonomy, the exploration cluster was further subdivided into Directed and Random exploration based on combinations of rolling accuracy and switch rate, capturing distinct modes of uncertainty-guided and value-independent sampling, respectively. Post-reversal trials, defined as the first five trials following a contingency change, were classified as a period of “Adaptation” since this period represents the greatest period of Q-value violations. The speed of adaptation was based on whether the animal achieved greater than chance accuracy during this period (fast: accuracy ≥ 0.6; slow: accuracy < 0.6), yielding seven states in total: Exploitation, Directed Exploration, Random Exploration, Left Spatial Bias, Right Spatial Bias, Fast Adaptation, and Slow Adaptations.

**Figure 7.**
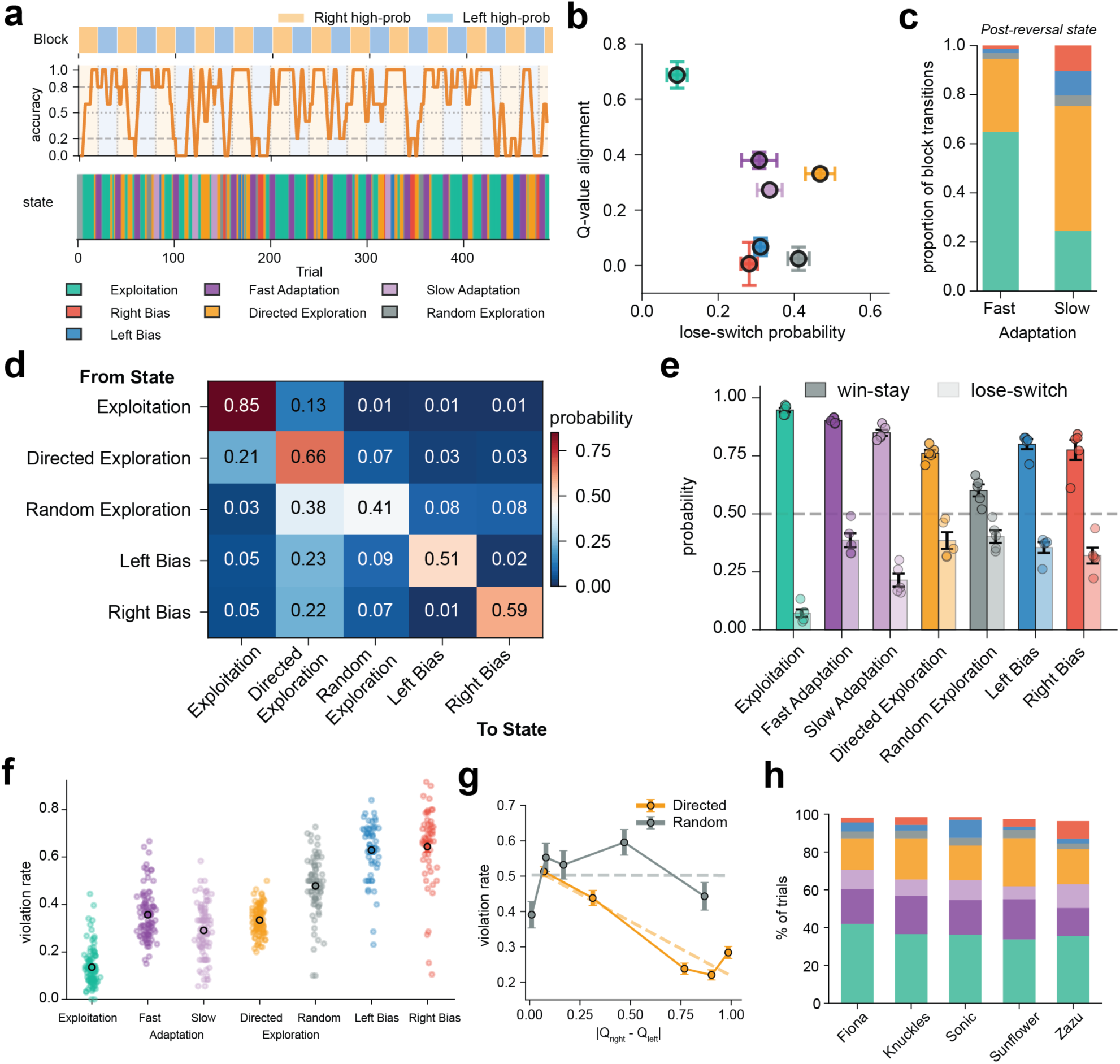
A behavioral state framework reveals distinct modes of reward-guided decision-making in marmosets. **a**, Representative sessions from a single animal (Knuckles) showing block structure (top), rolling accuracy computed over a five-trial window (middle), and trial-by-trial behavioral state classification (bottom). Colors correspond to the seven states defined in the legend. **b**, Behavioral state positioned in the space defined by lose-switch probability (x-axis) and Q-value alignment (y-axis), derived from the Q-learning model with choice perseveration term. Error bars indicate standard error of mean in both directions. **c**, State composition in the first trial following a fast versus slow block reversal. Colors indicate post-reversal behavioral state identity. Fast and slow refer to the speed of adaptation following a block reversal, classified within either the Adapting (fast) or Adapting (slow) state. **d**, Trial-to-trial state transition probability matrix, computed from all consecutive trial pairs pooled across animals and sessions and row-normalized to give the conditional probability of transitioning from each state (rows) to each state (columns). Diagonal entries reflect self-transition probability. **e**, Probability of repeating after a win (win-stay) or switching after a loss (loss-switch) by state classification averaged across all individual marmosets (dots). Dashed horizontal line indicates chance (0.5). **f**, Violation rate by state, defined as the proportion of trials on which animals chose the option with the lower model-estimated Q-value, restricted to trials where Q-values differed. Individual session-level estimates are shown as colored dots; black outlined circles indicate the animal mean **g**, Violation rate versus the Q-value difference fit with a linear regression line for both directed (yellow) and random (grey) exploration. **h**, State occupancy (% of trials) per behavioral state for each individual marmoset, pooled across all sessions.

To characterize the computational strategies associated with each state, we examined their positions in a two-dimensional space defined by lose-switch probability and Q-value alignment (**Figure 7b**). Exploitation trials had the lowest lose-switch probability and the highest Q-value alignment of all states (lose-switch: 0.092±0.027, alignment: 0.688±0.048), consistent with stable value-guided exploitation during late-block periods with high accuracy and low switch rates. Directed Exploration and Random Exploration trials differed markedly in Q-value alignment (0.331±0.017 and 0.024±0.042, respectively) but had similar lose-switch probabilities (0.468±0.039 and 0.411±0.028), indicating that these states were distinguished primarily by whether choices remained coupled to learned value rather than by sensitivity to negative feedback. Left Bias and Right Bias trials had low lose-switch probabilities and Q-value alignment near zero (alignment: 0.067±0.032 and 0.006±0.078, respectively), consistent with spatial preferences in these states being driven by positional habit rather than learned value. Fast and Slow Adaptation trials occupied intermediate positions in this space, with Fast Adaptation characterized by lower Q-value alignment than Slow Adaptation (0.241±0.025 versus 0.437±0.017), indicating that rapid adaptation required animals to choose the newly correct option before their value estimates had updated sufficiently to favor it. Lose-switch probability was numerically higher for fast than Slow Adaptation (0.387±0.037 versus 0.280±0.033) but did not differ significantly between groups.

To characterize how animals transition between states, we evaluated transition probabilities and performance measures across states (**Figure 7c, Supplementary Figure 4d**). In block transitions during which animals quickly adapted to the new contingency (Fast Adaptation), animals rapidly identified the new high-probability option (accuracy: 0.796±0.011) and most often transitioned directly into Exploitation, characterized by high accuracy (0.940±0.011) and low switch rates (0.057±0.010). In block transitions during which adaptation was slower (Slow adaptation), animals reached substantially lower accuracy over the same five trial post-reversal period (accuracy: 0.274±0.021) and rarely transitioned directly into Exploitation, instead entering Directed Exploration characterized by elevated switch rates (0.306±0.022) and moderate accuracy (0.636±0.014).

Beyond block transitions, we examined how animals transition out of each state (**Figure 7d**). Exploitation was the most self-sustaining state, with animals remaining in it on consecutive trials with a probability of 0.852±0.005. Directed Exploration was the most common transition into Exploitation from any non-Adaptation state (0.215±0.005), and Random Exploration transitioned most often into Directed Exploration (0.381±0.019) rather than directly into Exploitation, suggesting a sequential resolution in which value-independent exploration precedes uncertainty-guided exploration before exploitation resumes. Like Random Exploration, Left and Right Bias were less self-sustaining (0.508±0.027 and 0.586±0.023, respectively) and exited most often into Directed Exploration rather than Exploitation directly (Left Bias: 0.228±0.018; Right Bias: 0.220±0.018). Thus, while Fast Adaptation periods reached Exploitation directly, recovery from all other states proceeded through Directed Exploration as an intermediate.

While lose-switch vs the q-value alignment disentangled Exploitation from non-exploit states, the relationship between win-stay and lose-switch probabilities further revealed behavioral mechanisms underlying state distinctions (**Figure 7e**). Exploitation had the highest win-stay probability (0.950±0.008) and the lowest lose-switch probability (0.092±0.027), consistent with stable reward-guided exploitation under probabilistic feedback. Across both exploration and adaptation states, win-stay probability was the primary dissociation between groups. Fast adaptation had significantly higher win-stay than Slow (0.914±0.005 versus 0.892±0.003; U=23.0, p=0.032), while lose-switch did not differ significantly between these periods (0.387±0.037 versus 0.280±0.033; U=20.0, p=0.151), indicating that sensitivity to positive feedback rather than negative feedback distinguished rapid from slow post-reversal adaptation. Similarly, Directed Exploration had significantly higher win-stay than Random Exploration (0.828±0.009 versus 0.707±0.030; U=25.0, p=0.008), while lose-switch probabilities did not differ significantly (0.468±0.039 versus 0.411±0.028; U=18.0, p=0.310), indicating that sensitivity to positive feedback distinguished uncertainty-guided from value-independent exploration. Together, these results reveal a double dissociation in which win-stay probability tracks the degree of value-guided choice across states while lose-switch probability reflects a more uniform sensitivity to negative feedback that does not reliably differentiate behavioral modes.

Violation rates differed markedly across states and further distinguished the behavioral properties of each (**Figure 7f**). Exploitation trials showed the lowest violation rate of all states (0.138±0.016), consistent with animals reliably choosing the higher-valued option during stable exploitation. Fast adaptation trials showed higher violation rates than Slow adaptation periods (0.304±0.013 versus 0.378±0.010), consistent with fast learners choosing against their current Q-values to identify the newly correct option while slow learners followed their value estimates more faithfully during the post-reversal window. Directed Exploration showed intermediate violation rates (0.341±0.012) while Random Exploration showed substantially higher rates (0.501±0.017), consistent with the value sensitivity of Directed Exploration choices. Left Bias and Right Bias showed the highest violation rates of all states (0.633±0.019 and 0.654±0.030), confirming that spatial preferences in these states were largely dissociated from learned value.

To further distinguish Directed from Random Exploration, and to address the concern that state differences might reflect task-generated reward sequences rather than genuine differences in behavioral strategy, we examined violation rate as a function of Q-value difference, a measure that controls for local task history by asking whether animals in different states respond differently to equivalent value signals (**Figure 7g**). In Directed Exploration, violation rate decreased significantly as Q-value difference increased (slope=−0.301±0.027, p<0.0001), indicating that animals in this state were sensitive to the relative value of the two options and chose more reliably as the value signal strengthened. In Random Exploration, violation rate was insensitive to Q-value difference (slope=0.127±0.120, p=0.299), indicating that choices in this state were not guided by learned value regardless of how discriminable the options were. The slopes differed significantly between states (t=−5.300, p<0.0001), confirming that Directed and Random Exploration are mechanistically distinct: animals in Directed Exploration sample the environment in a value-sensitive manner while animals in Random Exploration do not. Importantly, Directed Exploration was the dominant form of exploration across all five animals (7731 versus 1455 trials), indicating that value-sensitive sampling was the primary exploratory mode across sessions and individuals (**Figure 7h**). The consistency of this state structure across all five animals suggests it reflects a general framework for reward-guided decision making rather than idiosyncratic individual strategies, raising the question of whether the same behavioral states emerge when reward feedback is made fully deterministic.

### Deterministic reward contingencies enhance value-based control through increased choice determinism

To test whether the reliability of reward feedback shapes decision making strategy, we compared behavior between probabilistic (80-20) and deterministic (100-0) reward contingencies in the five animals that performed both task variants (Fiona, Knuckles, Sonic, Sunflower, Zazu). Deterministic contingency trials were drawn from sessions in which animals performed a visual discrimination task with cued and uncued contingency changes (**Figure 2**), where pair of blocks containing an uncued reversal was included in the analysis.

Animals showed higher asymptotic accuracy under deterministic contingencies (80-20: 0.734±0.018, 100-0: 0.786±0.026; paired t-test: t(4)= −4.214, p=0.014; **Figure 8a**), and overall switch rate was lower outside of block transitions (80-20: 0.175±0.016, 100-0: 0.157±0.034; t(4)=3.464, p=0.026; **Figure 8b**), consistent with more stable choice behavior under deterministic feedback.

**Figure 8.**
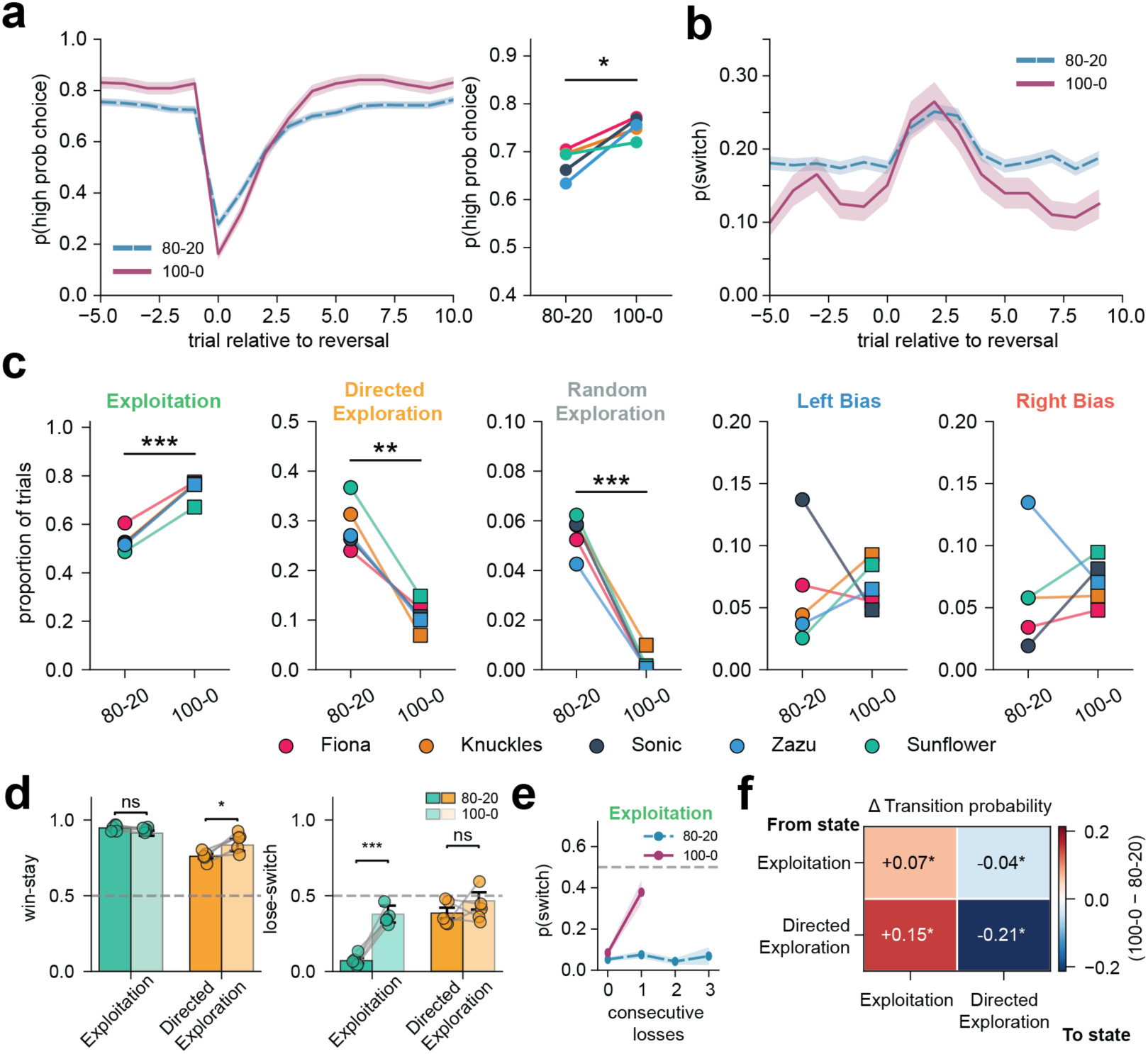
Deterministic reward contingencies reorganize behavioral strategy through state-specific changes in reward sensitivity. **a**, Probability of selecting the high-probability choice aligned to block reversal (trial position = 0) for probabilistic (80-20, blue, dashed) and deterministic (100-0, purple, solid) conditions (left). Shaded regions indicate ± SEM across animals. (right) Paired plot depicting the mean session accuracy per animal across conditions. *p<0.05, paired t-test (n = 5). **b**, Mean probability of switching (p(switch)) aligned to block reversal for 80-20 (blue, dashed) and 100-0 (purple, solid) conditions. Shaded regions indicate ±SEM. **c**, Proportion of trials spent in each behavioral state (excluding Adapting trials) for 80-20 and 100-0 conditions. Each panel shows a single state; circles and squares indicate individual animal means for 80-20 and 100-0 respectively, connected by lines for paired animals (n=5). **p<0.01, ***p<0.001, paired t-test. **d**, Win-stay (left) and lose-switch (right) probabilities for Exploitation and Directed Exploration states in 80-20 and 100-0 conditions. Bars indicate group means; circles indicate individual animal means for overlapping animals (n=5), connected by lines across conditions. *p<0.05, ***p<0.001, paired t-test. **e**, Probability of switching as a function of consecutive losses within the Exploitation state for 80-20 (blue, dashed) and 100-0 (purple, solid) conditions. Points indicate animal means; error bars indicate ±SEM. **f**, Change in trial-to-trial state transition probability between deterministic and probabilistic reward contingencies (100-0 minus 80-20), restricted to transitions between Exploitation and Directed Exploration states. Asterisks indicate significant differences between conditions (paired t-test, *p<0.05, n=5 animals).

To determine whether improved performance under deterministic feedback reflected a reorganization of behavioral strategy rather than uniform enhancement across states, we applied the trial-level classification framework to both conditions (**Figure 8c**). Exploitation occupancy increased under deterministic contingencies (80-20: 0.369±0.014, 100-0: 0.521±0.016; t(4)= −7.778, p=0.002), while Directed Exploration decreased (80-20: 0.202±0.016, 100-0: 0.076±0.009; t(4)=8.605, p=0.001) and Random Exploration was nearly absent (80-20: 0.038±0.002, 100-0: 0.002±0.001; t(4)=15.004, p < 0.001). Left Bias and Right Bias state occupancy did not differ between conditions, consistent with spatial habit formation being insensitive to reward reliability. The reduction in overall switch rate under deterministic feedback therefore reflects a shift in the distribution of behavioral states rather than a uniform suppression of switching across all modes of decision making.

To determine whether deterministic feedback altered reward sensitivity within behavioral states, we examined win-stay and lose-switch probabilities for Exploitation and Directed Exploration trials separately in each condition (**Figure 8d**). Win-stay probability in Exploitation was stable across conditions (80-20: 0.948±0.009, 100-0: 0.938±0.017; t(4)=0.995, p=0.376), indicating that animals were equally likely to repeat a rewarded choice regardless of feedback reliability. In contrast, lose-switch probability in Exploitation increased under deterministic contingencies (80-20: 0.072±0.017, 100-0: 0.364±0.056; t(4)=−9.622, p=0.001), indicating that animals were far more likely to switch following a loss when feedback was deterministic. In Directed Exploration, win-stay probability increased under deterministic contingencies (80-20: 0.761±0.015, 100-0: 0.857±0.040; t(4)=−3.641, p=0.022), while lose-switch probability was stable (80-20: 0.385±0.036, 100-0: 0.428±0.058; t(4)=−0.648, p=0.552). These results reveal a double dissociation: deterministic feedback selectively increased loss sensitivity within Exploitation and win sensitivity within Directed Exploration, demonstrating that reward reliability reshapes reward sensitivity in a state-specific and outcome-specific manner.

The increase in lose-switch probability within the Exploitation under deterministic feedback raised the question of whether animals were responding to individual losses or accumulating evidence across consecutive losses before switching. To address this, we examined switch probability as a function of consecutive losses within the Exploitation state separately for each condition (**Figure 8e**). Under probabilistic contingencies, switch probability was low and stable across loss streaks (loss streak 0: 0.053±0.009, loss streak 1: 0.075±0.018, loss streak 2: 0.043±0.015), consistent with animals in Exploitation largely discounting losses under probabilistic feedback regardless of how many occurred consecutively. Under deterministic contingencies, switch probability after zero consecutive losses was similar to the probabilistic condition (0.084±0.016) but increased dramatically after a single loss (0.378±0.056), after which consecutive loss trials were nearly absent within the Exploitation. The near-complete absence of lose-stay trials within Exploitation under deterministic feedback (Percent of lose-stay trials: 0.6%, deterministic vs. 15.3%, probabilistic) confirms that losses during exploitation triggered immediate switching rather than being discounted as noise. This asymmetry indicates that under deterministic feedback a single loss during exploitation functions as an unambiguous contingency change signal that triggers immediate error correction, while under probabilistic feedback losses during exploitation are treated as noise and discounted regardless of their frequency.

These findings demonstrate that reward reliability does not uniformly enhance decision making but selectively reshapes it. Deterministic feedback increased exploitation occupancy, reduced exploration, and sharpened error correction within exploitation, primarily by altering how animals respond to outcomes within states rather than by changing which states exist. Consistent with this, transition probabilities between Exploitation and Directed Exploration shifted coherently under deterministic feedback: Exploitation became more self-sustaining and Directed Exploration routed more rapidly back to Exploitation (**Figure 8f**), indicating that deterministic feedback accelerates recovery of exploitative control following exploratory excursions. The reduction in overall switching under deterministic feedback therefore reflects a reorganization of behavioral strategy rather than a suppression of flexibility.

## Discussion

Using a voluntary home-cage touchscreen task, we developed a curriculum-based training of marmosets from a simple visual discrimination task to a probabilistic 2-armed bandit behavior that provides a dynamic environment for probing decision making. In doing so, we show that adaptive decision making in marmosets is organized into discrete and persistent behavioral states whose structure and reward sensitivity are shaped by environmental predictability. Animals engaged with the task on their own initiative, retained memory of rewarded actions across bouts of disengagement, and adapted to contingency changes based on reward history alone without explicit cues or water restriction. A Q-learning model with choice perseveration captured trial-by-trial dynamics and comparison of observed choice consistency against the policy-predicted following rate indicated that violations of value-guided choice are consistent with the model’s stochastic policy rather than independent exploratory processes. A trial-level behavioral state classification framework identified distinct modes of decision making that differed systematically in their reliance on reward information and their transition structure. Finally, comparing behavior under probabilistic and deterministic reward contingencies revealed that reward reliability did not uniformly enhance performance but selectively reorganized state occupancy and reshaped reward sensitivity in a state-specific and outcome-specific manner.

### Engagement and disengagement: naturalistic testing reveals motivation as a variable

Unlike conventional testing paradigms that suppress motivational variability through water restriction or food deprivation, voluntary home-cage testing preserves the natural fluctuations in engagement providing a window into how motivation shapes decision making (Wallace et al., 2015; Vanderlip et al., 2026). Periods of inactivity occurred more frequently following unrewarded trials, indicating that negative feedback modulated not only choice updating but also the willingness to continue engaging with the task. This is consistent with accounts of motivation as an integral component of the decision making process rather than a separate modulatory influence (Niv et al., 2007; Kurzban et al., 2013; Guitart-Masip et al., 2014). Despite periods of disengagement, marmosets’ performance upon re-engagement reflected the contingency in effect before the pause rather than a reset to chance. This raises the question of whether marmosets retain the future rewarded action or a stronger memory of the unrewarded action that triggered disengagement. Nonetheless, periods of disengagement from the task may themselves support reward memory consolidation, as hippocampal sharp-wave ripples occurring during seconds-long periods of awake immobility have been shown to consolidate reward-associated experiences and facilitate subsequent goal-directed behavior (Jadhav et al., 2012; Papale et al., 2016).

### Behavioral state framework: what observable behavior reveals beyond RL parameters

The Q-value analysis revealed that violations of value-guided choice are consistent with the model’s stochastic policy but could not distinguish animals actively exploring to resolve that uncertainty from those perseverating on a spatial preference independent of learned value. A trial-level classification framework based entirely on observable choice metrics resolved this ambiguity by identifying distinct and persistent behavioral states that differed systematically in their reliance on reward information, their sensitivity to outcome history, and their transition structure. The persistence and structured transitions of these states suggest that marmoset decision making moves between epochs with fundamentally different computation or heuristics, consistent with hidden Markov model accounts of behavioral variability across rodents, non-human primates, and humans (Ebitz et al., 2019; Ashwood et al., 2022; Laurie et al., 2024). Transitions back to Exploitation from non-Adaptation states were rarely direct, with animals instead passing through Directed Exploration as an intermediate, suggesting that uncertainty-guided sampling may serve to destabilize entrenched behavioral policies before stable exploitation resumes. This interpretation carries testable circuit-level predictions, as distinct neural regions differentially track relative and total uncertainty during decision making (Daw et al., 2006; Badre et al., 2012). Directed and Random Exploration have further been shown to rely on dissociable neural substrates, providing a framework for identifying the circuits that govern transitions between behavioral states (Zajkowski et al., 2017; Tomov et al., 2020).

### Tipping the balance between exploration and exploitation

Comparing behavior under probabilistic and deterministic reward contingencies revealed that reward reliability did not uniformly enhance decision making but selectively reorganized it. Examining reward sensitivity within states revealed a double dissociation: deterministic feedback selectively increased loss sensitivity in Exploitation while increasing win sensitivity in Directed Exploration, demonstrating that reward reliability reshapes reward processing in a state-specific and outcome-specific manner rather than producing a uniform enhancement of reward sensitivity. Although consistent with normative accounts of adaptive learning in which the reliability of reward feedback determines how strongly individual outcomes update behavioral policy (Behrens et al., 2007; Nassar et al., 2010), our findings extend these accounts by revealing that reward predictability does not scale a single global learning rate but instead reconfigures the computational policy implemented within each behavioral mode independently.

The most striking manifestation of this reorganization was the response to single losses in the Exploitation state. Whereas consecutive losses during probabilistic feedback produced no systematic increase in switching, a single loss under deterministic feedback was sufficient to trigger immediate switching. This asymmetry reflects the optimal behavioral strategy whereas under deterministic contingencies, a single loss constitutes unambiguous evidence of a contingency reversal, making immediate switching the correct response. This is consistent with normative accounts in which the predictability of the environment determines the degree to which individual outcomes should update behavioral policy, with surprising outcomes carrying maximal informational weight precisely when the environment is otherwise reliable (Pearce and Hall, 1980; Nassar et al., 2010; McGuire et al., 2014).

### Circuit predictions grounded in current findings

The behavioral state transitions and reward sensitivity changes identified here generate specific and testable predictions about the neural circuits supporting adaptive control in marmosets. The structured transition pathway through Directed Exploration suggests a role for anterior cingulate cortex in signaling the need for updated strategies and gating the transition back to exploitation, consistent with its established role in tracking environmental volatility and regulating value updating (Behrens et al., 2007; Rushworth and Behrens, 2008; Kennerley et al., 2011). The persistence of Bias states and their resolution through Directed Exploration rather than direct transition to Exploitation suggests a role for orbitofrontal cortex, given its established role in restoring sensitivity to reward value following contingency changes and breaking habitual response tendencies (Clarke et al., 2008; Noonan et al., 2010). The state-specific increase in loss sensitivity under deterministic feedback suggests a role for dopaminergic signaling in ventral striatum (O’Doherty et al., 2004; Pessiglione et al., 2006), consistent with the idea that prediction error signals scale with the informational value of negative feedback rather than its raw frequency (Diederen et al., 2016; Dabney et al., 2020; Moller et al., 2022). The stability of Bias states across reward conditions, and their insensitivity to feedback reliability, implicates dorsolateral striatum in maintaining spatial habits independent of value-based control (Jackson et al., 2019). These predictions can be directly tested using the chronic recording and pharmacological manipulation capabilities of the home-cage platform described here.

### Marmosets as a laboratory model in modern neuroscience

Marmosets have emerged as a powerful model for studying the circuits that support flexible decision making across aging and disease. First, marmosets are a non-human primate with a lifespan that is amenable to studying development and aging. Namely, they have the shortest gestation of all primates (∼143 days), each birth results in twins or triplets and their lifespan is relatively short amongst primates (aged condition after 8-9 years). Second, an array of cognitive behaviors has established age-related changes in task performance and upon disease relevant perturbations (Glavis-Bloom et al., 2022; Vanderlip et al., 2023; Vanderlip et al., 2024). Third, causal manipulations of prefrontal and basal ganglia subregions in the marmoset, including orbitofrontal cortex, anterior cingulate cortex, dorsolateral prefrontal cortex and the striatum, produce specific and dissociable deficits in reversal learning and value-based choice (Roberts et al., 1994; Dias et al., 1996; Clarke et al., 2004; Clarke et al., 2005; Clarke et al., 2008; Rygula et al., 2010; Jackson et al., 2016; Jackson et al., 2019; Duan et al., 2021a). Finally, marmosets are amenable to viral vector-based manipulations, advanced imaging techniques and transgenic strategies, making them uniquely positioned to bridge the gap between the circuit-level mechanistic work possible in rodents and the cognitive complexity of human and other non-human primate behavior (Macdougall et al., 2016; Ebina et al., 2018; Vormstein-Schneider et al., 2020; Goertsen et al., 2022; Chen et al., 2025).

Together, these findings establish a framework for understanding how reward predictability shapes the balance between exploitation and exploration in a non-human primate. By combining voluntary home-cage behavioral testing with computational modeling and state-level analysis, we reveal that adaptive decision making is not a single process but an orchestration of distinct behavioral modes whose structure and reward sensitivity are shaped by the statistics of the environment. This platform and framework position the marmoset as a powerful model for dissecting the circuit mechanisms of flexible decision making across the lifespan and in the context of neuropsychiatric and neurodegenerative disease.

## Supplementary Figures

**Supplementary Figure 1.**
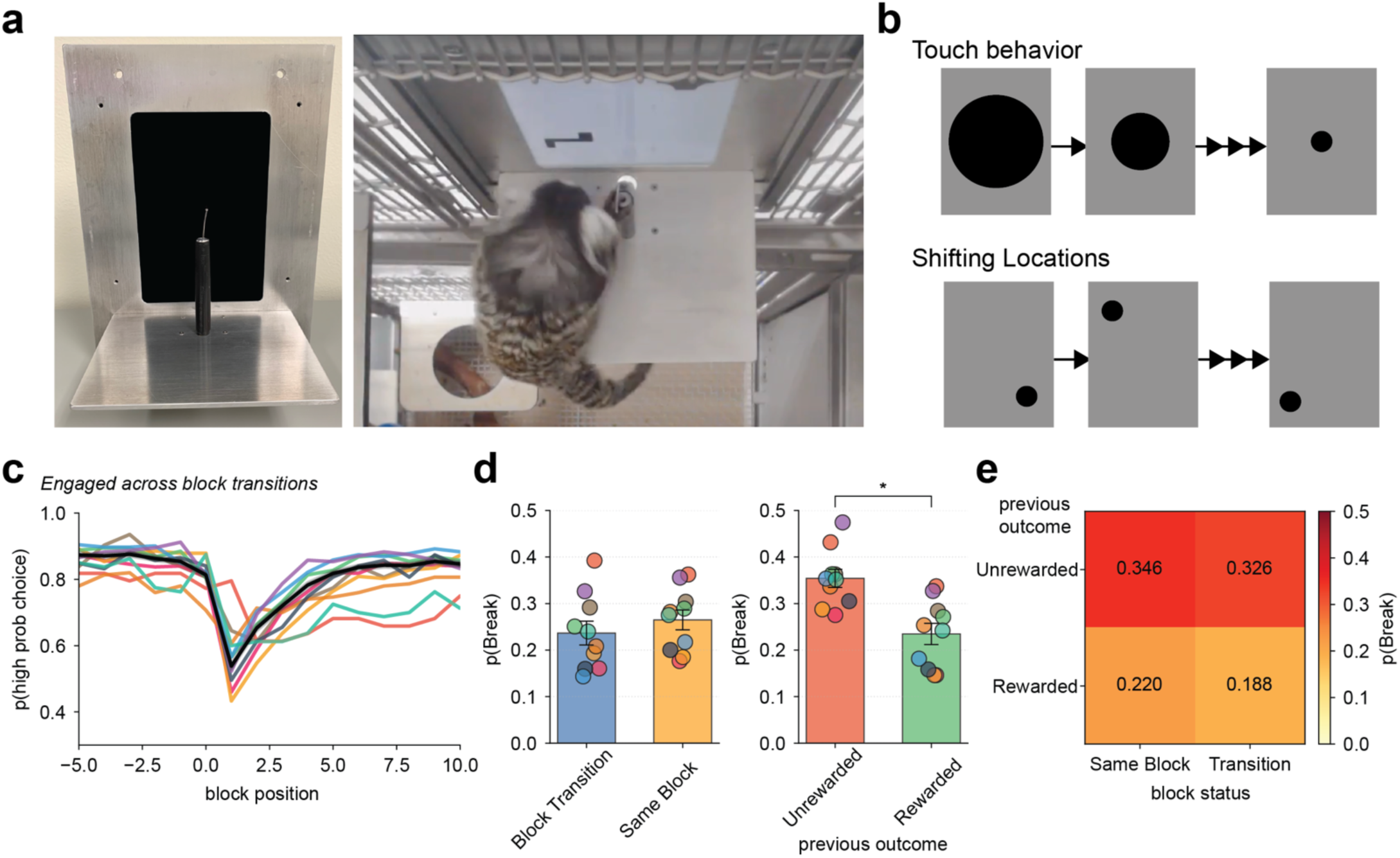
Tracking reward-based decision making, task acquisition and engagement utilizing a home-cage touchscreen. **a**, Picture of task screen holder device (left) and while attached to the home-cage (right) with a platform for marmosets to position in front of the screen and a port that provides sweetened liquid rewards **b**, Schematic representation of acquisition sequences. “Touch behavior” rewards any touch behavior within the black circle with decreasing size to enhance touch precision. Following successful touch behavior, marmosets interact with “shifting locations” where the smallest stimuli is moved to all possible positions on the screen and rewarded for all touches within stimuli. **c**, Probability of selecting the high probability choice (‘high-prob choice’) at each trial position for each individual marmoset (colored lines) or the average (black line). **d**, Probability of a disengagement (p(break)) following a block transition or within the same block (left) or after a rewarded or unrewarded outcome (right) across all individual marmosets (dots). **e**, Heatmap representing the relationship between the previous outcome and block status on the probability of taking a break

**Supplementary Figure 2.**
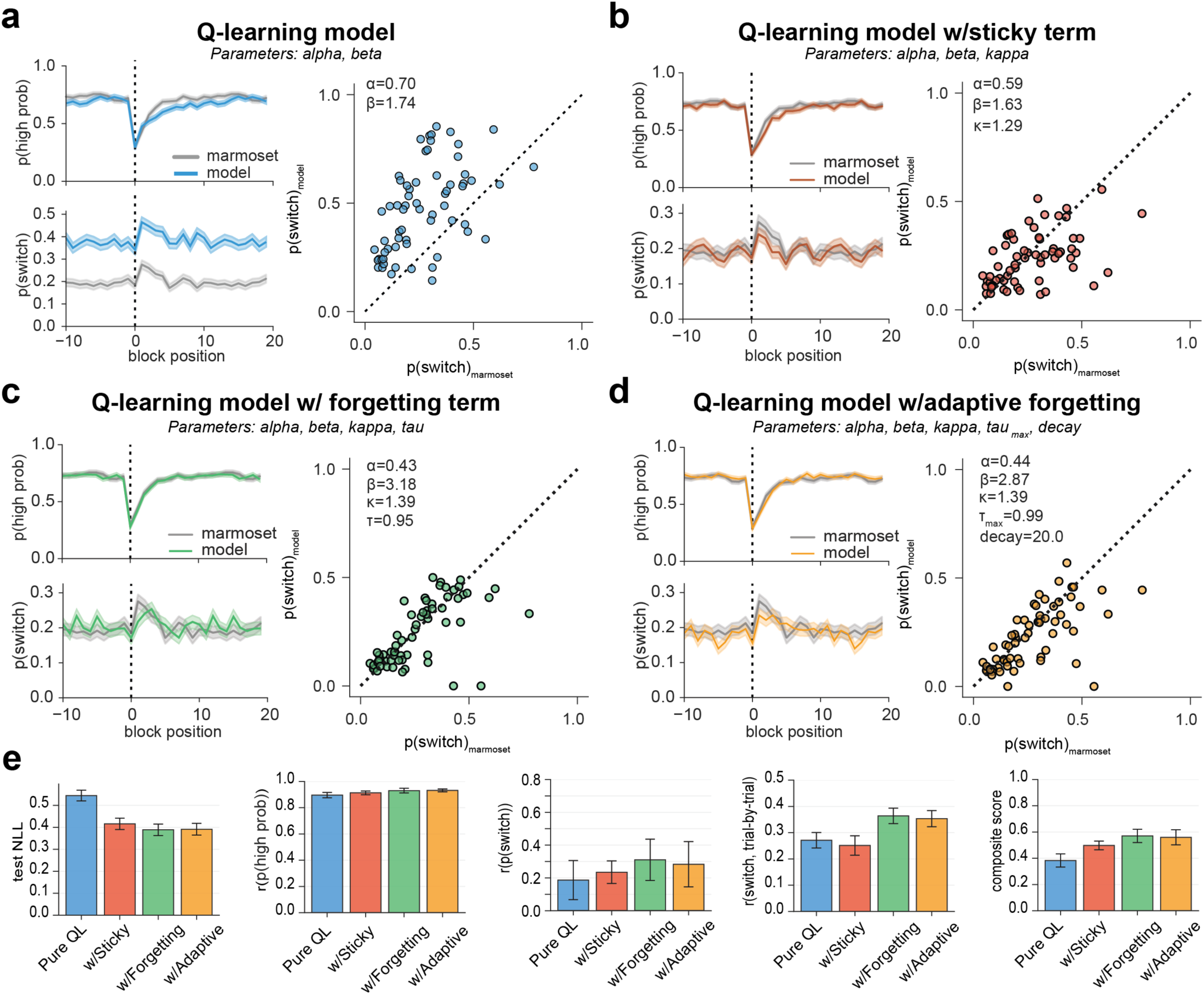
Q-learning model with choice kernel captures trial-by-trial dynamics of the 2-armed bandit performance and captures differences across individuals. **a**, Pure Q-learning model (parameters: α, β). Mean p(high probability choice) (top) and p(switch) (bottom) aligned to block reversal (trial position = 0, dashed vertical line) for marmosets (grey) and model simulations (blue). Shaded regions indicate ±SEM. Right: Session-level p(switch) predicted by the model versus observed marmoset p(switch). Each dot represents one session. Dashed line indicates unity. Mean fitted parameters shown. **b**, As shown in panel (**a**) but for the Q-learning model with sticky term (parameters: α, β, κ). **c**, As shown in panel (**a**) but for the Q-learning model with forgetting term (parameters: α, β, κ, τ). **d**, As shown in panel (**a**) but for the Q-learning model with adaptive forgetting (parameters: α, β, κ, τ_max_, decay). **e**, Model comparison across four metrics: test negative log-likelihood (NLL), correlation between predicted and observed p(high probability choice), correlation between predicted and observed p(switch), trial-by-trial correlation of switching behavior, and composite score averaging across all metrics. Bars indicate mean ±SEM across sessions.

**Supplementary Figure 3.**
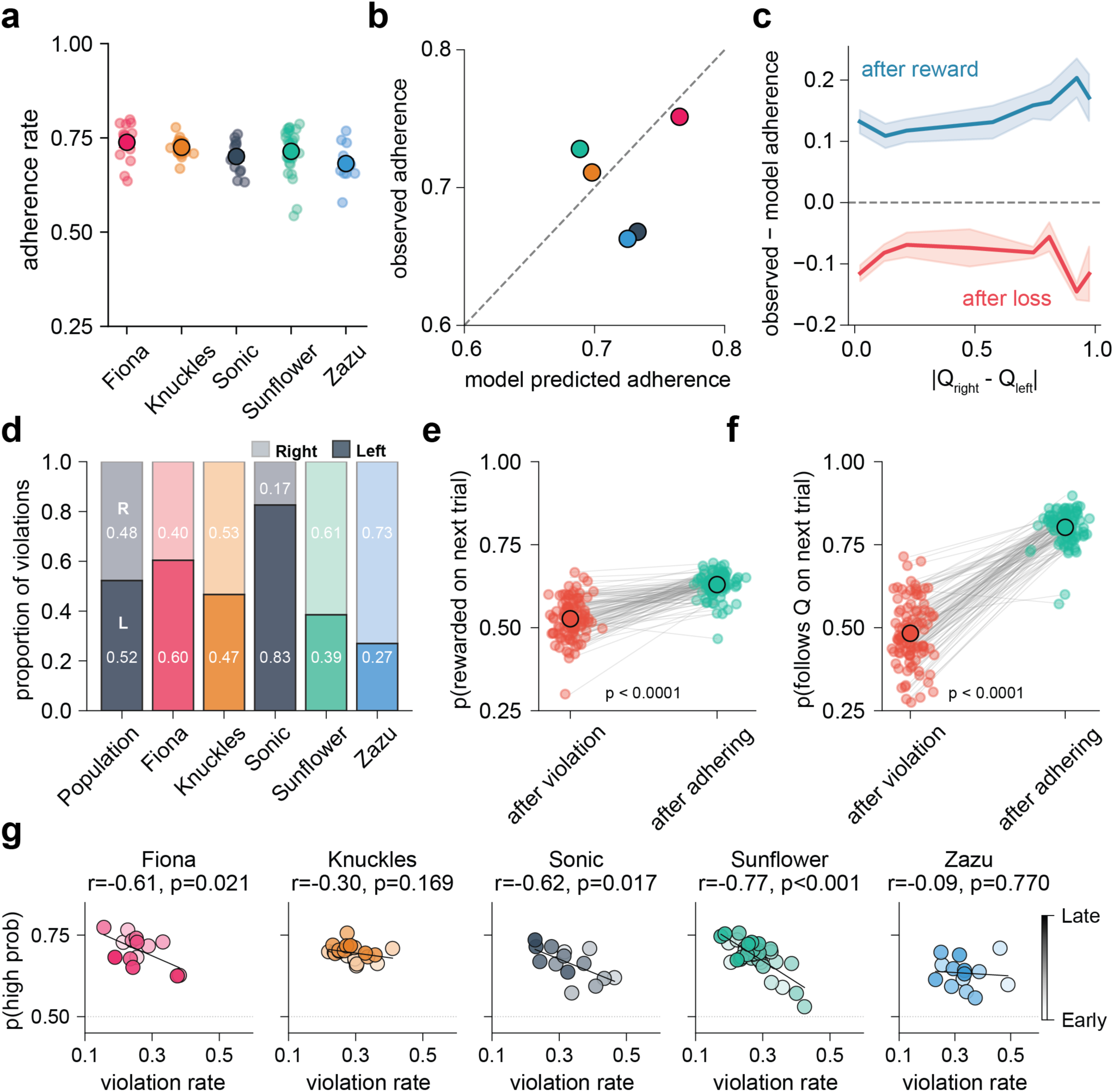
Q-value following behavior across animals, block position, outcome history, and session performance. **a**, Probability of adhering to Q-values (p(follows Q-values) by animal. Each dot represents one session; large filled circles indicate the animal mean. **b**, Observed probability of adhering to Q-values as a function of the policy-predicted adherence rate. Each dot represents one animal; the dashed line indicates the identity. Animals scatter symmetrically around the identity with no systematic bias (t(4)=−0.871, p=0.433). **c**, Difference between observed and model-predicted probability of following as a function of absolute difference between Q_right_ and Q_left_. Lines show animal means following a rewarded (blue) or unrewarded (red) outcome; shaded regions indicate ± SEM. **d**, Direction of Q-value violations by animal. Stacked bars show the proportion of violations directed toward the left (L, opaque) versus right (R, partially transparent) side. **e**, Probability of being rewarded on the next trial following a violation versus a following trial. Each dot is one session; gray lines connect matched sessions. Open circles show the mean across all animals ± SEM (p < 0.0001, paired t-test). **f**, Probability of following Q-values on the trial after a violation versus a following trial. Violations are followed by reduced Q-value adherence on the subsequent trial (p < 0.0001, paired t-test). **g**, Session-level violation rate versus accuracy (p(High Prob)) for each animal. Each dot represents one session, colored light to dark by session date within animal (light = early, dark = late). Dashed lines show linear regression fits. Violation rate was negatively correlated with session accuracy for Fiona (r=-0.61, p=0.021), Sonic (r=-0.62, p=0.017), and Sunflower (r=-0.77, p<0.001), but not for Knuckles (r=-0.30, p=0.169) or Zazu (r=-0.09, p=0.770).

**Supplementary Figure 4.**
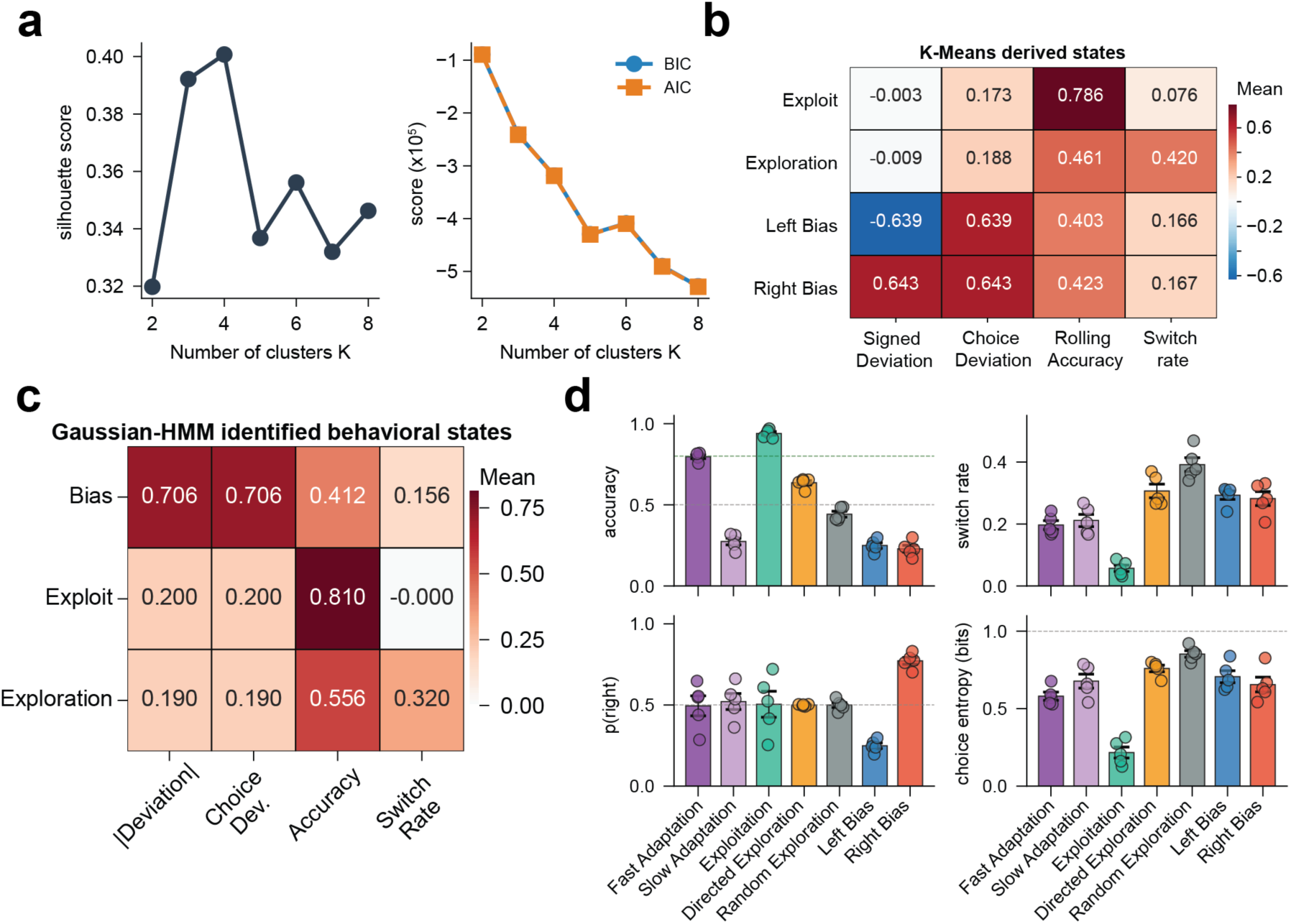
Data-driven validation of the behavioral state taxonomy. **a**, Silhouette score (left) and Bayesian Information Criterion (BIC) and Akaike Information Criterion (AIC) scores (right) across K-means cluster solutions from K=2 to K=8. **b**, Mean feature values for each of four behavioral clusters identified by K-Means clustering (K=4) applied to derived behavioral features without reference to state labels. Signed Deviation, the signed difference between an animal’s choice probability and the expected choice probability given block structure, served as the primary feature distinguishing bias from non-bias states (blue box). Cluster profiles were computed from train-set centroids and evaluated on held-out sessions. **c**, HMM emission means for each state in the three-state solution, confirming the behavioral signatures of Bias (high absolute deviation, low accuracy), Exploitation (high accuracy, near-zero switching), and Exploration (moderate accuracy, high switching), derived independently of the K-Means solution. **d**, Behavioral metrics by state classification across seven behavioral states (Adapting fast, Adapting slow, Exploitation, Directed Exploration, Random Exploration, Left Bias, Right Bias). (top, left) probability of selecting the high probability choice, (top, right) switch rate, (bottom, left) probability of selecting the rightward option, (bottom, right) choice entropy (bits). Bars indicate the mean across all animals; dots represent individual session averages. **a-c.** N=5 animals, n=100 sessions total. All classification and clustering analyses were performed on held-out sessions not used for threshold derivation.

## Methods

All procedures were performed in accordance with the guidelines established by the National Institutes of Health and were approved by the Broad Institute at MIT and Harvard Institutional Animal Care and Use Committee.

### Marmosets

Common marmosets (*Callithrix jacchus*) of both sexes were used for all marmoset behavioral experiments. Marmosets were housed in socially enriched environments and maintained under standard colony conditions. All behavioral procedures were conducted during the light phase of the housing cycle and were completed within the home-cage with visual, tactile and auditory access to cage-mates at all times. No water regulation was required.

### Behavioral Apparatus

Marmoset experiments were conducted within the animals’ home-cage environment, where the behavioral device was positioned in one quadrant of the cage and the marmoset of interest was allowed access for 1-3 hours per session. The apparatus consisted of a custom-designed attachment integrated with the caging system to present a touchscreen-based operant setup. A reward delivery port was positioned on a vertical pole approximately 2 inches from the screen to spatially separate stimulus selection from reward consumption. Liquid rewards consisted of diluted apple juice or diluted Ensure. Reward delivery was calibrated on a per-session basis, and total sugar intake was controlled to a maximum of 10 g per session across all conditions. Task control, stimulus presentation, response detection, and reward delivery were implemented using MonkeyLogic, a MATLAB-based behavioral control framework developed by the National Institute of Mental Health, interfaced with an Arduino-controlled pump and display hardware.

### Touchscreen Behavioral Acquisition

Prior to task-specific training, marmosets underwent a touchscreen behavioral acquisition phase to establish reliable interaction with the touchscreen and association between touch responses and reward delivery. Animals were first habituated to the presence of the touchscreen apparatus within the home-cage environment and allowed to freely explore the device during initial sessions. During early acquisition, touches anywhere on the screen were reinforced to promote engagement with the touchscreen. Over successive sessions, reinforcement was gradually restricted to touches directed toward visual stimuli to shape stimulus-specific responding. The size and contrast of visual stimuli were adjusted as needed to facilitate acquisition until the visual stimuli were the same size that would be employed in the task design (**Supplementary Figure 1B**). Finally, the stimuli were moved to all possible screen positions to ensure that the marmoset was successfully tracking object location (**Supplementary Figure 1B**). Once animals reliably initiated touch responses to visual stimuli, reward delivery was contingent upon correct stimulus selection according to task rules. Acquisition was considered complete when animals consistently engaged with the touchscreen, demonstrated stable touch accuracy, and completed enough trials per session to support task-specific training. Touchscreen acquisition sessions were conducted during the light phase, and animals were allowed to engage with the apparatus for up to 1-3 hours per session. Progression through acquisition stages was individualized based on performance to ensure consistent touchscreen interaction prior to formal task training.

### Task Design

Three touchscreen-based behavioral tasks were used to assess cognitive flexibility and reward-based decision making. All tasks employed the same touchscreen interface and reward delivery system described above, and animals interacted with visual stimuli via direct touch responses. Each trial consists of two choice stimuli that appear on the screen and upon selection will either provide a liquid reward or no feedback. While the stimuli presentations differ across the three tasks, they share a common set of reward contingencies including both deterministic (100/0) and probabilistic (80/20) schedules. For each task, the rule is kept constant for a series of trials, known as a “block”. To maximize reward acquisition, animals must acquire the rule within the block and identify the stimuli associated with higher reward probability and avoid stimuli associated with lower reward probability. As described below, these tasks represent structured variations of a two-armed bandit framework designed to probe decision-making across conditions.

1. <u>Visual Discrimination Task</u> In the visual discrimination task, animals were presented with two distinct visual stimuli that differed in shape and/or appearance. On each trial, the touchscreen was divided into four quadrants, and the two stimuli appeared simultaneously in two of the four possible positions. Stimulus locations varied pseudo-randomly across blocks such that the spatial position of each stimulus changed from block to block. One stimulus was consistently rewarded, while the other was unrewarded. Animals were required to identify and select the rewarded stimulus dependent of its spatial location.
2. <u>Reversal Learning Task:</u> In the reversal learning task, the same two visual stimuli were presented simultaneously at two fixed locations on the touchscreen. Within a given block of trials, one stimulus was designated as the highly rewarded stimulus. At an uncued point within the block, reward contingencies reversed such that the previously unrewarded or lowly rewarded stimulus became highly rewarded, and the previously rewarded stimulus became unrewarded or lowly rewarded. This task required animals to update stimulus-reward associations following changes in contingency.
3. <u>Two-Armed Bandit Task:</u> In the two-armed bandit task, two identical visual stimuli were presented at fixed positions on the touchscreen. Stimulus identity and spatial location remained constant across trials and sessions. Reward outcomes were delivered probabilistically, with reward probabilities assigned independently to each stimulus. At uncued points, reward contingencies reversed, requiring animals to update action-outcome associations and adjust their choice behavior accordingly.

### Task Progression

Following touchscreen behavioral acquisition, marmosets were exposed to a variety of task designs across their lifespan. For most subjects, marmosets were first trained on the visual discrimination task, followed by the reversal learning task, and then the two-armed bandit task. This progression was designed to establish basic stimulus-reward associations before introducing increasing levels of contingency uncertainty. For each task and reward contingency condition, animals were trained until a minimum of 1,000 completed trials was obtained and was not dependent on accuracy or task performance. Achieving this criterion typically required 5-7 sessions per condition, depending on individual engagement and performance. Periodically, marmosets underwent prolonged periods without any touchscreen exposure. Upon re-entry into the training progress, marmosets were often progressed through shaping before entering into one of the three tasks.

#### Behavioral Analysis

All behavioral analyses were performed on completed trials in which animals made a valid choice between the two available options. Trials in which no response was registered or in which task requirements were not met were excluded from analysis. Behavioral metrics were computed using custom scripts written in MATLAB and Python.

To characterize task performance, behavior was aligned to block structure. Within each task, reward contingencies were held constant within blocks of trials until either a cued transition or an uncued reversal. Block position was defined relative to the point of contingency change, with block position 0 corresponding to the first trial following the change. Behavioral metrics were quantified as a function of block position to assess how animals adapted their choices following changes in stimulus and/or reward contingencies.

Choice behavior was summarized using multiple complementary metrics. Performance was quantified as the probability of selecting the higher-rewarded option (p(high prob choice)) as a function of block position. To capture behavioral flexibility, the probability of switching choices between consecutive trials (p(switch)) was also computed. These metrics allowed dissociation of reward tracking from exploratory or perseverative choice strategies. Additionally, a series of conditional probabilities were employed throughout the paper. This includes when the probability an animal stays following a rewarded outcome (win-stay) or switches following a loss (lose-switch).

All analyses were performed using the same computational pipeline for both species, with task-specific parameters adjusted as described in the *Task Design* sections.

#### Computational Modeling

All computational models were fit to either individual choice sequences or the observable features outlined within each model at the session level using L-BFGS-B optimization with 50 random initializations per session. Model performance was evaluated on held-out data using 80/20 train-test splits stratified by block. Sessions in which any parameter reached its optimization bound were flagged as boundary cases and excluded from parameter-level comparisons.

### Reinforcement Learning models

A family of reinforcement learning models was fit to individual choice sequences from the 80-20 two-armed bandit task. All models maintained action value estimates (Q-values) for the left and right choice options, updated on each trial according to a prediction error:

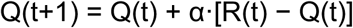

where α is the learning rate (0 < α < 1) and R(t) is the binary reward outcome (1 for rewarded, 0 for unrewarded). Q-values were initialized to 0.5 at the start of each session. Choice probability was determined by a softmax policy:

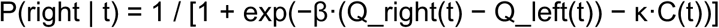

where β is the inverse temperature governing choice determinism, κ is the choice perseveration parameter, and C(t) is the choice kernel tracking whether the animal repeated (C(t) = 1) or switched (C(t) = −1) relative to the previous trial.

Four model variants were compared. The base model (pure Q-learning) included α and β only. The choice kernel model added κ as a free parameter. The asymmetric learning rate model replaced α with separate parameters for positive (α+) and negative (α−) prediction errors. The forgetting model added a decay parameter that reduced the value of the unchosen option toward 0.5 on each trial.

Models were fit by minimizing negative log-likelihood using L-BFGS-B optimization with 50 random initializations per session to avoid local minima. Parameter bounds were α ∈ [0.0001, 0.9999], β ∈ [0.001, 50], and κ ∈ [−5, 10]. Model comparison used the Bayesian Information Criterion (BIC), with lower values indicating better fit penalized for model complexity. Sessions in which any parameter reached its optimization bound were flagged as boundary cases and excluded from parameter-level comparisons. Model fit was assessed on held-out test trials using 80/20 train-test splits stratified by block.

### Model Selection

As stated in the results section, model selection was based on the ability for the model to predict trial-by-trial performance as depicted in the following analyses: probability of selecting the high-prob choice across block transitions, probability of switching across block positions and the comparison between conditional switching probabilities for the marmoset vs the model (**Supplementary Figure 2a-d**). The Q-learning model with choice kernel substantially outperformed a model with no choice history (pure Q-learning; ΔBIC=+546.45±101.93 across all animals; **Supplementary Figure 2a-b**). A model with separate learning rates for positive and negative prediction errors did not improve fit over the kernel model (ΔBIC=+109.18±30.10 relative to forgetting model; **Supplementary Figure 2c**). A model incorporating forgetting of the unchosen option’s value had the lowest absolute BIC, but three of five animals had forgetting parameters at the upper bound of the parameter space, indicating parameter instability **(Supplementary Figure 2d**). The kernel model was therefore selected as the working model for subsequent analyses based on parameter stability and interpretability. The kernel model captured trial-by-trial fluctuations in choice behavior, correctly predicting choices on 81.0±1.4% of held-out test trials and accurately reproducing the temporal dynamics of log-odds of choosing right vs left (**Supplementary Figure 2e**).

The choice kernel model was selected as the working model based on superior BIC relative to pure Q-learning (ΔBIC = +546.45 ± 101.93) and parameter stability relative to the forgetting model, in which three of five animals showed forgetting parameters at the upper bound of the parameter space. Trial-by-trial Q-value estimates from the selected model were used in subsequent analyses of Q-value following, violation rate, and behavioral state validation.

### Behavioral state classification

A trial-level behavioral state classifier was developed to assign each trial to one of seven mutually exclusive states based on observable choice metrics computed over a five-trial sliding window: rolling accuracy (proportion of trials selecting the high-probability option), rolling switch rate (proportion of trials in which the animal switched from the previous choice), and spatial choice deviation (absolute difference between the animal’s rolling choice probability and the expected choice probability given block structure). All metrics were derived from observable behavior without reference to the reinforcement learning model.

The classifier applied a hierarchical set of rules based on thresholds derived from K-Means cluster midpoints identified in the validation analysis (*see Validation methods below*). Trials occurring within the first five trials following a contingency reversal were assigned to the Adaptation state, defined by block position rather than behavioral metrics, reflecting the post-reversal updating period during which Q-values had not yet converged to the new contingency. Among remaining trials, spatial bias states were identified first: trials in which absolute spatial deviation exceeded 0.4 were assigned to Left Bias or Right Bias depending on whether the animal’s rolling choice probability toward the non-rewarded side exceeded 0.6. Among non-biased trials, Exploitation was assigned when rolling accuracy was at or above 0.6 and rolling switch rate was at or below 0.2, reflecting stable high-accuracy exploitation with minimal switching. Directed Exploration was assigned when rolling switch rate was at or above 0.2 and rolling accuracy was at or below 0.6, reflecting active uncertainty-guided sampling. Random Exploration was assigned when rolling accuracy fell below 0.4, reflecting value-independent stochastic choice. Trials not meeting any of these criteria were assigned to Exploitation as the default late-block state.

Adaptation trials were subdivided into Fast or Slow based on whether the animal reached above-chance accuracy (p(high-probability choice) >= 0.6) by the end of the five-trial post-reversal window. Directed and Random Exploration were distinguished by the criterion above rather than by Q-value information, consistent with the finding that these states could not be separated by unsupervised clustering of behavioral metrics alone but were distinguishable by their differential sensitivity to Q-value difference as confirmed in held-out analyses.

State transition probabilities were computed at the session level by counting consecutive trial-pair transitions within each session and normalizing each row to sum to one, then averaging normalized matrices across sessions to give equal weight to each session regardless of trial count. State dwell times were computed as the mean number of consecutive trials spent in each state before a transition. Win-stay and lose-switch probabilities were computed per animal by identifying trials preceded by a rewarded or unrewarded outcome respectively, then computing the proportion of stay or switch responses, and averaging across sessions within each animal before computing group means and standard errors. Q-value alignment was computed as the proportion of trials on which the animal chose the option with the higher model-estimated Q-value, restricted to trials where the absolute Q-value difference exceeded zero. Choice entropy was computed as the binary entropy of the rolling choice probability within each trial’s five-trial window.

### Validation of Behavioral State Identification

To identify whether distinct behavioral states could be recovered from observable choice metrics without reference to the reinforcement learning model, we applied K-Means clustering and a Gaussian Hidden Markov Model (HMM) to four derived behavioral features: absolute deviation from the optimal choice probability (|Rolling_PRight − Expected_PRight|), spatial choice deviation, rolling accuracy, and rolling switch rate, each computed over a five-trial sliding window. Sessions were split into training (80%) and held-out test (20%) sets at the session level prior to any model fitting, and all reported metrics reflect performance on held-out sessions.

1. **K-Means clustering.** To determine the natural number of behavioral clusters, K-Means clustering was applied to the training set across a range of solutions (K = 2–8). The optimal number of clusters was selected using the silhouette coefficient, which peaked at K = 4, corresponding to states aligned to Exploitation, Exploration, Left bias, and Right bias. Features were standardized exclusively on the training set and applied without refitting to the test set. The final K=4 solution was fit with 20 random initializations and the solution with lowest inertia retained. Cluster labels were matched to behavioral state names using the Hungarian algorithm to maximize overlap between cluster assignments and manual state labels on the training set. Agreement between K-Means clusters and manual labels on held-out sessions was quantified using the adjusted Rand index (ARI = 0.58).
2. **Gaussian Hidden Markov Model.** A Gaussian HMM with diagonal covariance was fit to the same four behavioral features to capture the sequential structure of state transitions across trials. The number of states was set to K = 3, corresponding to Bias, Exploitation, and Exploration, following model selection using log-likelihood, BIC, and AIC across K = 2–7. Absolute deviation was used in place of signed deviation to prevent the model from splitting left and right spatial bias into separate states based on choice direction rather than behavioral dynamics; left and right bias were instead assigned post-hoc by combining the HMM Bias state with block context (given by High_Prob_Is_Right, where Right block is 1 and left block is 0). Features were standardized using a scaler fit on training sessions only. To prevent singular covariance estimates arising from the discrete six-value lattice structure of rolling window features, a small amount of Gaussian noise (σ = 0.01) was added to scaled features prior to fitting, which was confirmed not to affect state assignments. The first five trials of each session were excluded to avoid contamination of rolling window features by the preceding session. Session boundaries were explicitly provided to the model via the lengths parameter in hmmlearn, preventing spurious state transitions across session boundaries. K-Means centroids derived from the training set were used to initialize HMM emission means, with transition matrices and covariances initialized randomly. Ten random restarts were run for up to 200 EM iterations each, and the solution with the highest training log-likelihood was retained. State sequences were decoded using the Viterbi algorithm. Agreement between HMM-decoded states and manual labels was quantified on held-out test sessions (3-state ARI = 0.487; 4-state post-hoc ARI = 0.474), with a train/test gap of 0.006 confirming no overfitting. State dwell times were derived analytically from self-transition probabilities as 1/(1 − p_ii).
3. **Supervised validation.** To assess whether behavioral states were recoverable from raw trial-by-trial observables without access to the rolling window features used in state definition, a Random Forest classifier (300 trees, maximum depth 8, balanced class weights) was trained on choice history (t−1 to t−3), reward history (t−1 to t−3), and win-stay/lose-shift indicators, using the same session-level train/test split. Overall classification accuracy on held-out sessions was 73% without block context. The addition of a binary indicator of which side carried the higher reward probability increased accuracy to 82%, confirming that behavioral states are recoverable from raw behavior and that the 16-percentage-point gain from block context reflects the inherent ambiguity between correct spatial persistence and spatial bias in raw choice history alone.

#### Statistics

All data are reported as mean ± SEM unless otherwise noted. Statistical comparisons were performed in Python using SciPy. For comparisons against chance performance, Wilcoxon signed-rank tests were used. For within-animal comparisons across conditions, paired t-tests were used with animals as the unit of observation. For session-level correlations between violation rate and accuracy, Pearson correlation coefficients were computed per animal. Slopes relating violation rate to Q-value difference were estimated by linear regression per state, with significance assessed by t-test on the regression coefficient.

Where multiple comparisons were performed simultaneously, including comparisons of state proportions across conditions (**Figure 7c**) and win-stay and lose-switch probabilities across states and conditions (**Figure 7d**); p-values were corrected for multiple comparisons using the Benjamini-Hochberg false discovery rate procedure with a threshold of q < 0.05. Uncorrected p-values are reported for individual planned comparisons.

Significance thresholds are indicated as follows, unless otherwise noted: *p < 0.05, **p < 0.01, ***p < 0.001.

State transition probabilities were compared across conditions using session-level averages in which each session contributed equally regardless of trial count. Effect sizes for paired comparisons are not reported given the small sample size of five animals, which precludes reliable effect size estimation.

## Data availability

All behavioral data, analysis code, and model fitting scripts will be made publicly available upon publication at the following GitHub link: https://github.com/mastrok/marmoset_behav.

